# Glycosaminoglycan-mediated lipoprotein uptake protects cancer cells from ferroptosis

**DOI:** 10.1101/2024.05.13.593939

**Authors:** Dylan Calhoon, Lingjie Sang, Divya Bezwada, Nathaniel Kim, Amrita Basu, Sheng-Chieh Hsu, Anastasia Pimentel, Bailey Brooks, Konnor La, Ana Paulina Serrano, Daniel L. Cassidy, Ling Cai, Vanina Toffessi-Tcheuyap, Vitaly Margulis, Feng Cai, James Brugarolas, Ryan J. Weiss, Ralph J. DeBerardinis, Kıvanç Birsoy, Javier Garcia-Bermudez

## Abstract

Lipids are essential for tumours because of their structural, energetic, and signaling roles. While many cancer cells upregulate lipid synthesis, growing evidence suggests that tumours simultaneously intensify the uptake of circulating lipids carried by lipoproteins. Which mechanisms promote the uptake of extracellular lipids, and how this pool of lipids contributes to cancer progression, are poorly understood. Here, using functional genetic screens, we find that lipoprotein uptake confers resistance to lipid peroxidation and ferroptotic cell death. Lipoprotein supplementation robustly inhibits ferroptosis across numerous cancer types. Mechanistically, cancer cells take up lipoproteins through a pathway dependent on sulfated glycosaminoglycans (GAGs) linked to cell-surface proteoglycans. Tumour GAGs are a major determinant of the uptake of both low and high density lipoproteins. Impairment of glycosaminoglycan synthesis or acute degradation of surface GAGs decreases the uptake of lipoproteins, sensitizes cells to ferroptosis and reduces tumour growth in mice. We also find that human clear cell renal cell carcinomas, a distinctively lipid-rich tumour type, display elevated levels of lipoprotein-derived antioxidants and the GAG chondroitin sulfate than non-malignant human kidney. Altogether, our work identifies lipoprotein uptake as an essential anti-ferroptotic mechanism for cancer cells to overcome lipid oxidative stress in vivo, and reveals GAG biosynthesis as an unexpected mediator of this process.

---

Evidence since the early 1950s indicates that cancer cells activate de novo lipid synthesis to fuel their rapid growth^1,2^. Paradoxically, tumours intensify the uptake of lipids encapsulated within lipoproteins that circulate in the blood. This phenomenon has been observed in lymphoma^3^, kidney cancer^4^, melanoma^5^, and glioblastoma^6^, and raises the possibility that intracellularly synthesized lipids and lipids taken from circulation may fulfill different needs for cancer cells. The two major types of lipoproteins in humans are low density and high density lipoproteins (LDL and HDL, respectively). Some tumours increase import of LDL^3,6^ and HDL^4^, but the precise mechanisms enabling this exacerbated lipoprotein uptake are unknown.

Tumours experience oxidative stress during all stages of cancer progression due to the generation of reactive oxygen species (ROS). One major deleterious effect of ROS accumulation is lipid peroxidation, which disrupts membrane function and leads to a non-apoptotic and iron-dependent form of cell death called ferroptosis^7^. Susceptibility to lipid peroxidation has emerged as a potential avenue for new cancer therapies. Growing evidence suggests that lipid peroxides accumulate in tumours during radiation therapy^8^ and in cancer cells undergoing metastasis^9^, and that ferroptosis-suppressing mechanisms promote cancer progression. Cancer cells have developed two categories of mechanisms to escape ferroptosis: enzymatic systems that convert lipid peroxides into non-toxic metabolites^10^, and the accumulation of antioxidant metabolites that inhibit lipid peroxidation through diverse mechanisms. The latter category contains lipid antioxidants, such as lipophilic vitamins D^11^, E^12^ and K^13^, coenzyme Q10 (CoQ10)^14,15^, monounsaturated fatty acids (MUFAs)^16^, squalene^3^ and 7-hydroxycholesterol^11^.

It is unknown which mechanisms cancer cells employ to take up lipoproteins, what advantages are bestowed by enhanced lipoprotein uptake, and whether imported lipids serve different functions than lipids synthesized by the cell. Thus, deciphering the functional nodes involved in lipid uptake is critical to understanding the role of exogenously-acquired lipids in tumorigenesis.

## Lipoprotein supplementation renders cancer cells resistant to ferroptosis

To understand how lipoproteins support cancer cell survival and proliferation, we performed a pooled CRISPR genetic screen in HeLa cells (Fig. 1a), which avidly take up both LDL and HDL (Extended Data Fig. 1a, 1b). We transduced HeLa cells with a focused sgRNA library targeting 200 rate-limiting and cancer-relevant metabolic genes^3^, and passaged them in culture media with lipoprotein-depleted serum that either was or was not supplemented with physiologically-relevant levels of lipoproteins^17^ (Fig. 1a). Determination of sgRNA abundance and gene dependency revealed that the essentiality of most genes remained unchanged between the two conditions (Fig. 1b). Glutathione peroxidase 4 (*GPX4*) scored as the gene whose essentiality most significantly changed depending on lipoprotein availability: *GPX4* was essential in the absence of lipoproteins, but dispensable in lipoprotein-supplemented conditions (Fig. 1b, 1c, Extended Data Fig. 1c, 1d).

**Figure 1.**
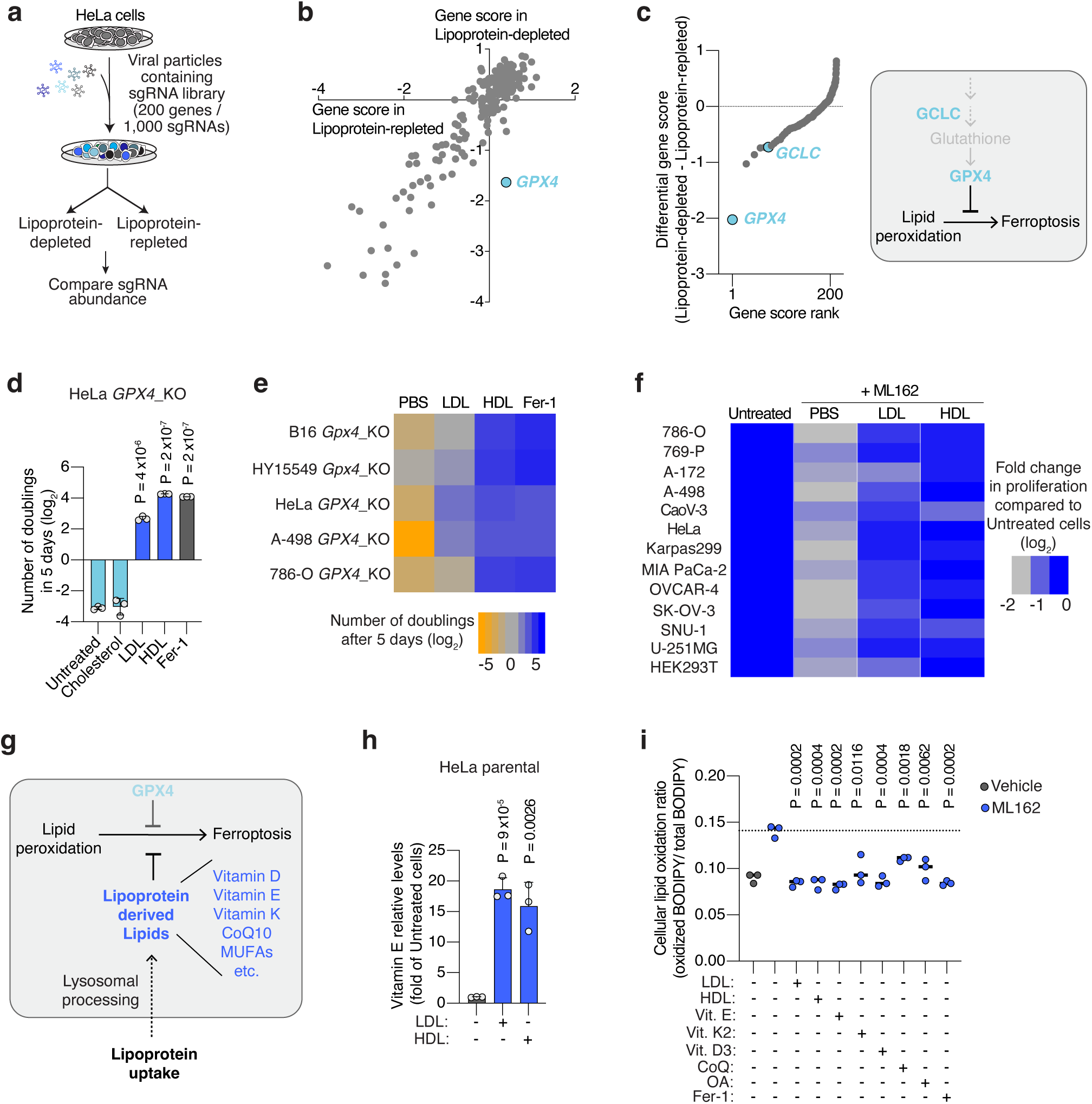
Lipoprotein supplementation promotes cancer cell resistance to ferroptosis. **a.** Scheme of CRISPR screen in HeLa cells transduced with a focused-metabolism sgRNA library and depleted of or supplemented with human lipoproteins. **b.** Gene essentiality graph showing changes in essentiality based on lipoprotein availability. *GPX4* becomes non-essential in cells supplemented with lipoproteins. **c.** Rank of genes whose essentiality in HeLa cells is most changed based on lipoprotein availability. Anti-ferroptotic genes are highlighted in blue (left). Scheme of the glutathione peroxidase pathway inhibiting lipid peroxidation and ferroptosis in cells (right). **d.** Number of doublings (log_2_) in 5 days of HeLa *GPX4*_KO cells in vitro either untreated or supplemented with cholesterol (5 μg/mL), human low density lipoproteins (LDL, 50 μg/mL), high density lipoproteins (HDL, 50 μg/mL) or ferrostatin-1 (Fer-1, 1 μM). **e.** Heatmap showing the number of doublings (log_2_) in 5 days of mouse (*Gpx4*) or human (*GPX4*) knockout cell lines in cell culture under the supplementation of PBS, LDL (50 μg/mL), HDL (50 μg/mL) or Fer-1 (1 μM). **f.** Heatmap showing the fold change in proliferation relative to untreated cells (log_2_) of a panel of lines with or without the GPX4 inhibitor ML162, and in the presence or absence of LDL (50 μg/mL) or HDL (50 μg/mL). **g.** Scheme showing the multiple anti-ferroptotic lipids that lipoproteins potentially carry, and that cancer cells can assimilate upon lipoprotein uptake and lysosomal processing. **h.** Mass spectrometry analysis of vitamin E levels in HeLa cells in the absence of lipoproteins or supplemented with LDL (50 μg/mL) or HDL (50 μg/mL). Data is presented as fold of metabolite levels in lipoprotein-depleted cells. **i.** Cellular lipid oxidation ratio using BODIPY-C11 in A-498 cells in the absence (gray) or presence (blue) of a GPX4 inhibitor (ML162, 250 nM) under supplementation or not of LDL (50 μg/mL), HDL (50 μg/mL), vitamin E (10 μM), vitamin K2 (10 μM), vitamin D3 (10 μM), CoQ10 (10 μM), OA (250 μM) or Fer-1 (1 μM). **d, h,** Bars represent mean ± s.d.; **i,** bars represent the median; **d-f, h, i,** n=3 biologically independent samples. Statistical significance determined by a two-tailed unpaired t-test compared to untreated cells (**d, h)** or ML162 treated cells (**i**).

GPX4 is the major enzyme that converts oxidized lipids into non-toxic metabolites in a glutathione-dependent manner and thus protects cells from ferroptosis^10^. Notably, the essentiality of another gene in this pathway was also among the most dependent on lipoprotein availability: glutamate-cysteine ligase catalytic subunit (*GCLC*), which encodes an enzyme required for glutathione synthesis (Fig. 1c, Extended Data Fig. 1c).

These genetic screen results suggested that supplementing lipoproteins might confer ferroptosis resistance in cancer cells lacking GPX4. To test this possibility, we used CRISPR to reduce GPX4 expression in five cancer cell lines: two derived from mouse tumours (melanoma B16 and pancreatic ductal adenocarcinoma cell line HY15549) and three patient-derived human cancer cell lines (HeLa, and clear cell renal cell carcinoma cell lines A-498 and 786-O) (Extended Data Fig. 1e, 1f). In line with previous studies, GPX4-deficient cell lines could not survive or proliferate in standard culture medium containing 10% fetal bovine serum^7^ (Fig. 1d). Treating cells with the lipophilic radical scavenger and ferroptosis inhibitor ferrostatin-1 (Fer-1) improved cell survival and enabled proliferation (Fig. 1d). However, supplementation of GPX4-deficient cells with physiologically-relevant concentrations of LDL or HDL promoted survival and proliferation in all five cell lines (Fig. 1d, 1e, Extended Data Fig. 1g). HDL supplementation provided an even stronger anti-ferroptotic effect than LDL, whereas cholesterol, a major component of lipoproteins, did not rescue cells from ferroptosis (Fig. 1d, 1e, Extended Data Fig. 1g). These experiments show that a critical benefit of lipoprotein uptake is the inhibition of ferroptosis through a cholesterol-independent mechanism, and that the magnitude of this anti-ferroptotic effect varies among lipoprotein classes.

To further generalize these findings, we assessed proliferation of a collection of 12 patient-derived cancer cell lines originating from blood, kidney, pancreas, ovarian, gastric and brain tumours, under chemical inhibition of GPX4 and in the presence of LDL or HDL. While the magnitude of the effect of HDL was again higher than that of LDL in most cell lines, supplementation of either lipoprotein class significantly promoted survival and proliferation of all cancer cell lines under chemically-induced lipid peroxidation stress (Fig. 1f). We also observed a consistent anti-ferroptotic effect of lipoprotein supplementation in non-transformed HEK293T cells. Therefore, our data shows that lipoprotein supplementation robustly suppresses ferroptosis in cell lines derived from many tissue types, including multiple cancers.

At least five different lipid species carried by lipoproteins have anti-ferroptotic potential (Fig. 1g). Vitamins D, E and K, and CoQ10 can act as radical trapping antioxidants within lipid membranes. Moreover, phospholipids containing MUFAs, such as oleic acid (OA), render cell membranes resistant to lipid peroxidation due to their reduced susceptibility to undergo peroxidation^16^. In humans, CoQ10, MUFAs, and the precursor of vitamin D3, 7-dehydrocholesterol, can be synthesized de novo or obtained from the diet, while vitamins E and K are exclusively obtained from the diet. Vitamin E primarily travels in the bloodstream within lipoproteins^18^, with a minor fraction binding to albumin or specific transport proteins. To confirm the transport of these anti-ferroptotic lipids by lipoproteins in our system, we cultured HeLa cells in lipoprotein-depleted media supplemented with either LDL or HDL, and quantified cellular vitamin E levels using mass spectrometry. Consistent with the role of lipoproteins in transporting the majority of lipids in the bloodstream, cells supplemented with either lipoprotein class exhibited over a 15-fold increase in vitamin E levels compared to lipoprotein-depleted cells (Fig. 1h).

To determine which lipids within lipoproteins inhibit ferroptosis in cancer cells, we supplemented candidate lipids individually. First, we assessed proliferation of GPX4-deficient cells, and observed enhanced proliferation upon supplementation of free vitamins D3, E, K2, CoQ10, or albumin-conjugated OA in four different GPX4-deficient cell lines (Extended Data Fig. 2a, 2b). Second, we measured oxidized lipid levels during chemical GPX4 inhibition, and observed reduced lipid oxidation in lymphoma or clear cell renal cell carcinoma (ccRCC) cells upon supplementation of LDL, HDL, free vitamin D3, E, K2, CoQ10, or albumin-OA (Fig. 1i, Extended Data Fig. 2c, 2d). This confirms the protective effect of lipoprotein-free vitamin E, K2, CoQ10, or OA against ferroptosis, as reported previously.

Next, we aimed to assess the individual roles of lipids within lipoproteins in protecting against lipid peroxidation. Using a genetic approach, we knocked out genes required for the anti-ferroptotic function of several lipid species (Extended Data Fig. 2e). Specifically, we targeted Acyl-CoA Synthetase Long Chain Family Member 3 (*ACSL3*)^16^ and Apoptosis Inducing Factor Mitochondria Associated 2 (*AIFM2*), also known as Ferroptosis Suppressor Protein 1 (FSP1)^14,15^, in lymphoma and ccRCC cells (Extended Data Fig. 2f). ACSL3 catalyzes the first committed step of MUFA incorporation into phospholipids, whereas FSP1 converts oxidized forms of vitamin K2^13^ and CoQ10 to their reduced, lipid peroxide-quenching quinone. However, supplementation of lipoproteins reversed the lipid peroxidation increase induced by the GPX4 covalent inhibitor ML162 in cells with or without ACSL3 (Extended Data Fig. 2g) or AIFM2 (Extended Data Fig. 2h), suggesting that neither lipoprotein-MUFAs nor lipoprotein-derived CoQ10 and vitamin K2 solely account for the anti-ferroptotic effect of lipoprotein lipids. Due to the lack of reported enzymes or genes regulating the anti-ferroptotic effect of vitamins D3 and E, we were unable to employ similar genetic approaches to isolate their antioxidant contribution upon lipoprotein supplementation.

Our data indicates that lipoproteins may transport distinct lipids with redundant anti-ferroptotic effects. This raises the possibility that inhibiting the avid uptake of these potent, antioxidant-rich lipoproteins is necessary to promote tumour ferroptosis.

## Cancer glycosaminoglycans are required for lipoprotein uptake and ferroptosis resistance

We next defined molecular components of lipoprotein transport. We devised a dual genetic screen platform in Karpas299, a lymphoma cell line with high lipoprotein uptake^3^. After transducing Karpas299 cells with a sgRNA library targeting metabolic genes and transporters^19^ (30,000 sgRNAs targeting 3,000 genes), we performed two genetic screens. First, cells were subjected to a proliferation-based screen in the presence or absence of the GPX4 inhibitor, ML210 (Fig. 2a), to define genes essential for cancer cell resistance to ferroptosis. In parallel, transduced lymphoma cells were incubated with low density lipoproteins labeled with a fluorescent lipophilic dye (DiI-LDL), followed by a Fluorescence-Activated Cell Sorting (FACS) approach to isolate cells with the most (top 5% fluorescence) and least (bottom 5% fluorescence) DiI-LDL uptake (Fig. 2a). We then isolated genomic DNA from each population, sequenced sgRNA amplicons, and compared sgRNA abundance. Each screen revealed a set of canonical genes involved in ferroptosis resistance or lipoprotein uptake. The oxidoreductase *AIFM2* and Peroxiredoxin 6 (*PRDX6*)^20^ scored as essential in the GPX4 inhibition screen (Fig. 2b, Extended Data Fig. 3a, 3b). Conversely, two members of the Sterol regulatory element binding protein (SREBP) system, *SREBF2* and *SCA*P^21^, were among the most essential genes in the lipoprotein uptake screen (Fig. 2c, Extended Data Fig. 3c).

**Figure 2.**
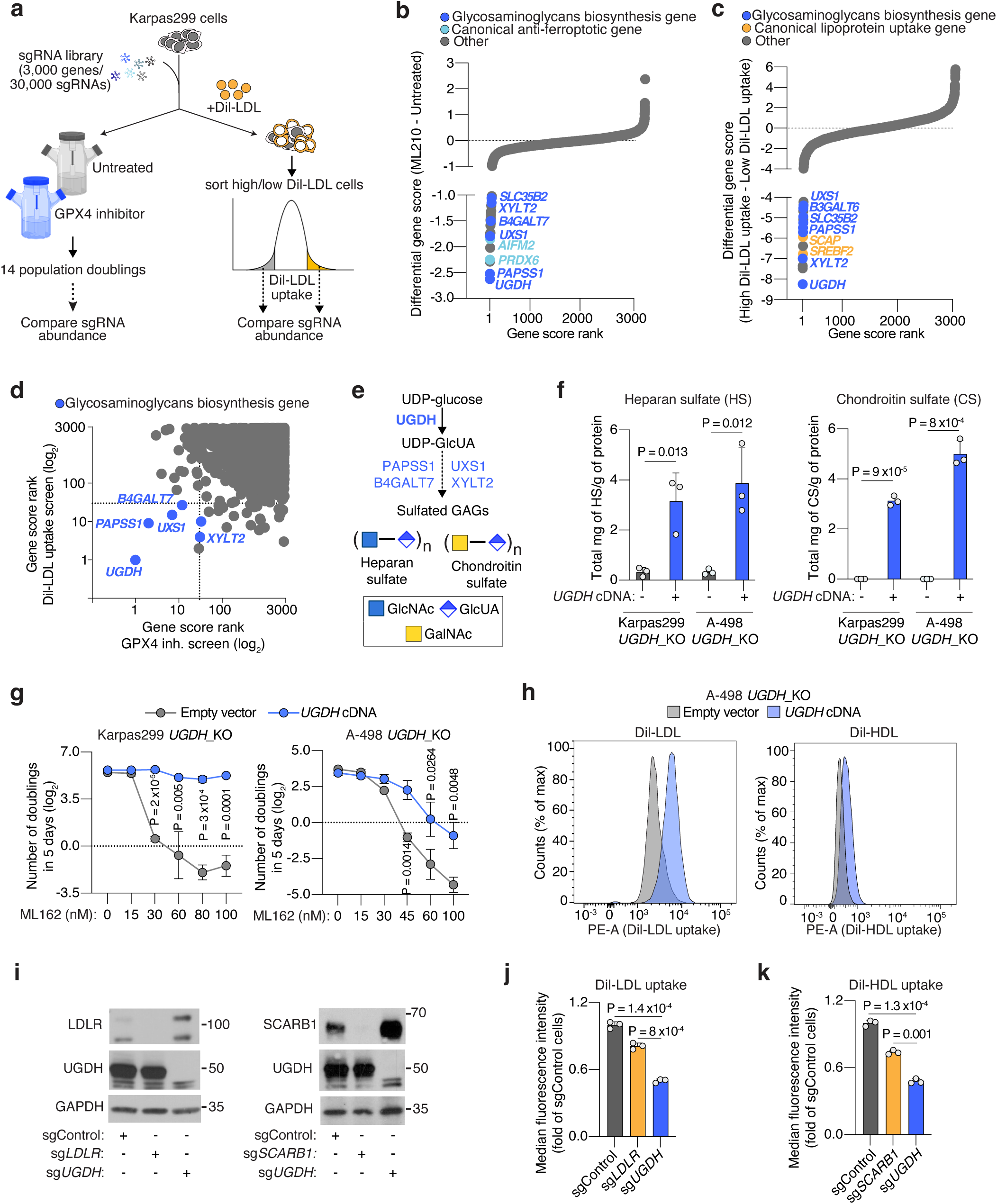
Cancer cells depend on the biosynthesis of glycosaminoglycans (GAGs) to take up lipoproteins, resist ferroptosis, and grow as tumours. **a.** Schematic of parallel CRISPR screens in Karpas299 lymphoma cells transduced with a metabolism-focused sgRNA library (3,000 genes). First, a proliferation-based screen in the presence or absence of a GPX4 inhibitor (ML210) during 14 population doublings (left); second, a FACS-based screen where cells were incubated with DiI-LDL for 2 hours and subjected to sorting for 5% highest and 5% lowest fluorescent cells (right). **b.** Rank of most essential genes under ML210 treatment compared to untreated Karpas299 cells. Canonical anti-ferroptotic genes are shown in light blue, and glycosaminoglycan biosynthesis genes are shown in dark blue. **c.** Rank of most essential genes for DiI-LDL uptake in Karpas299 cells. Lipoprotein uptake genes are shown in orange and glycosaminoglycan biosynthesis pathway genes are shown in dark blue. **d.** Gene scores ranks for the GPX4 inhibition and DiI-LDL uptake screens in Karpas299. All genes essential in both screens are part of the glycosaminoglycan biosynthesis pathway (dark blue). **e.** Glycosaminoglycans, such as heparan sulfate and chondroitin sulfate, are formed by disaccharides repeats containing glucuronic acid (GlcUA) and other monosaccharides. The most upstream and rate-limiting enzyme of this pathway is UGDH. All genes in the pathway that scored in CRISPR screens are highlighted in blue. **f.** Quantification of total milligrams of heparan sulfate (HS, left) and chondroitin sulfate (CS, right) per grams of protein in the indicated *UGDH*_KO cell lines expressing a sgRNA resistant *UGDH* cDNA (blue) or an empty vector (grey). **g.** Number of doublings in 5 days (log_2_) of the indicated *UGDH*_KO cell lines expressing a sgRNA resistant *UGDH* cDNA (blue) or an empty vector (grey) under the indicated concentrations of the GPX4 inhibitor, ML162. **h.** Representative flow cytometry plots showing the cellular uptake of DiI-LDL (left) or DiI-HDL (right) in A-498 *UGDH*_KO cells expressing a sgRNA resistant *UGDH* cDNA (blue) or an empty vector (grey). **i.** Immunoblot analysis of LDLR, SCARB1, and UGDH in HeLa cells transduced with a sgControl, sg*LDLR*, sg*SCARB1* or sg*UGDH*. GAPDH is included as a loading control. **j.** Fold change in the uptake of DiI-LDL of the indicated HeLa cell lines relative to sgControl-transduced cells assessed by flow cytometry. **k.** Fold change in the uptake of DiI-HDL of the indicated HeLa cell lines relative to sgControl-transduced cells assessed by flow cytometry. **f, g, j, k,** Bars or lines represent mean ± s.d.; **f, g, j, k,** n=3 biologically independent samples. Statistical significance determined by two-tailed unpaired t-tests.

Next, we focused our attention on essential genes for both DiI-LDL uptake and ferroptosis resistance (Fig. 2d), as we reasoned that this would pinpoint the pathways that tumours employ to take up antioxidant-rich lipoproteins. Remarkably, all the genes intersecting both processes in lymphoma cells belong to the same metabolic pathway: the biosynthesis of sulfated glycosaminoglycans (GAGs) (Fig. 2b-d, Extended Data Fig. 3a-c).

GAGs are long, linear polysaccharides consisting of repeating disaccharide units^22^. Their synthesis, assembly, and sulfation depend on multiple cytosolic or Golgi-localized enzymes, several of which scored in our screens (Fig. 2d, 2e). An almost universal component of the disaccharide repeats that constitute the two most abundant sulfated GAGs, heparan sulfate (HS)^23^ and chondroitin sulfate (CS)^24^, is glucuronic acid (GlcUA) (Fig. 2e). The enzyme responsible for GlcUA synthesis, uridine-diphospho glucose dehydrogenase (*UGDH*), was the top scoring gene for both resistance to GPX4 inhibition and increased DiI-LDL uptake (Extended Data Fig. 3b-e). We thus generated HeLa, mouse non-small lung cell carcinoma (Tom2), human renal cell carcinoma (A-498, Caki-2), and Karpas299 *UGDH*_KO cells (Extended Data Fig. 3f). We confirmed that *UGDH*_KO cells had negligible levels of HS and CS using mass spectrometry, whereas expression of a sgRNA-resistant *UGDH* cDNA increased levels of both GAGs (Fig. 2f, Extended Data Fig. 3f). Consistent with our screens, Karpas299, A-498, HeLa, Caki-2 and Tom2 UGDH-deficient cell lines were exquisitely more sensitive to GPX4 inhibition (Fig. 2g, Extended Data Fig. 3g, 3h), or glutathione synthesis inhibition mediated by erastin^25^ (Extended Data Fig. 3i) than their isogenic UGDH-expressing counterparts.

To our knowledge, GAG biosynthesis has not been linked to ferroptosis resistance. Sulfated GAGs are negatively charged at physiological pH, raising the possibility that they bind and chelate divalent metals^26^ like iron, an essential driver of the Fenton reaction that leads to ferroptosis. However, cells with no GAG biosynthesis displayed identical iron homeostasis and handling capacity compared to isogenic cells with GAGs (Extended Data Fig. 4a, 4b). Moreover, *UGDH*_KO cells did not have lower protein levels of the anti-ferroptotic effectors GPX4, cystine-uptake transporter SLC7A11^27^ or AIFM2/FSP1 than UGDH-expressing isogenic counterparts (Extended Data Fig. 4c). These results suggest that GAGs promote ferroptosis resistance through a mechanism independent of iron chelation, FSP1, and the GPX4-glutathione axis.

Prior studies have suggested that negatively-charged sulfated GAG chains can electrostatically bind and attract LDL due to positively-charged amino acids in the apolipoprotein fraction^28,29^. Therefore, we next tested whether GAGs render cancer cells resistant to ferroptosis by facilitating lipoprotein uptake. Indeed, lipoprotein uptake assays in HeLa, A-498, and Karpas299 cells revealed that UGDH-expressing cells had 2 to 3-fold higher DiI-LDL uptake (Fig. 2h, Extended Data Fig. 4d, 4e) and 2-fold higher DiI-HDL uptake (Fig. 2h, Extended Data Fig. 4f) than UGDH-deficient isogenic lines.

Non-transformed cells take up the majority of LDL and HDL through the LDL-receptor (LDLR)^30^ and the scavenger receptor class B1 (SCARB1)^31^, respectively. We aimed to define the contribution of GAGs to lipoprotein uptake in cancer cells relative to these canonical uptake pathways. We generated isogenic KO lines for *LDLR*, *SCARB1,* or *UGDH* in HeLa cells (Fig. 2i), and compared their LDL and HDL uptake capacity to those of cells transduced with a control sgRNA (sgControl). Knockout of *LDLR* or *SCARB1* decreased DiI-LDL or DiI-HDL uptake only mildly compared to those triggered by the loss of *UGDH* (∼50% reduction) (Fig. 2j, 2k). These experiments show that the GAG biosynthesis enzyme UGDH is a major determinant of LDL and HDL uptake in cancer cells.

The metabolic product of UGDH, uridine diphosphate glucuronic acid (UDP-GlcUA), has additional functions beyond serving as a building block for GAGs. To confirm that GAG biosynthesis is the pathway driving lipoprotein uptake and ferroptosis resistance in our model, we disrupted two additional GAG biosynthesis enzymes downstream of UGDH and tested whether these phenocopied *UGDH* loss. First, we tested the relevance of the sulfation of GAGs in ferroptosis resistance. 3’-Phosphoadenosine 5’-Phosphosulfate Synthase 1 (*PAPSS1*) is the enzyme that synthesizes 3’-phosphoadenylylsulfate (PAPS). Once PAPS is imported into the Golgi by the transporter SLC35B2, it is used by cellular sulfotransferases as a sulfate donor^32^ (Extended Data Fig. 5a). Both *PAPSS1* and *SLC35B2* are instrumental for GAG sulfation, and they scored in our screens (Fig. 2b-d, Extended Data Fig. 5b). Knocking out *PAPSS1* in Karpas299 cells (Extended Data Fig. 5c) decreased total levels of HS (Extended Data Fig. 5d), and reduced the number of sulfates per disaccharide unit in cellular HS to 30% of those from sgControl isogenic cells (Extended Data Fig. 5e). Lastly, when we subjected these cells to proliferation assays in the presence of a GPX4 inhibitor, we found that sg*PAPSS1* cells were much more sensitive to ferroptosis than sgControl cells (Extended Data Fig. 5f). We also tested the role of another GAG-forming monosaccharide: xylose. UDP-xylose is synthesized from UDP-GlcUA in the Golgi by the enzyme UDP-GlcUA decarboxylase 1 (UXS1), prior to its incorporation into a GAG chain^33^ by xylosyltransferase 2 (XYLT2) (Extended Data Fig. 5g). Both *UXS1* and *XYLT2* scored in the ferroptosis sensitivity and lipoprotein uptake screens (Fig. 2b-d, Extended Data Fig. 5h). Knocking out *UXS1* in Karpas299 cells (Extended Data Fig. 5i) triggered a 55% decrease in cellular HS levels (Extended Data Fig. 5j) and increased sensitivity to GXP4 inhibition (Extended Data Fig. 5k). Finally, we compared the lipoprotein uptake capacity of isogenic Karpas299 cell lines with an intact GAG biosynthesis capacity to those lacking the ability to synthesize UDP-GlcUA (sg*UGDH*), PAPS (sg*PAPSS1*), or UDP-xylose (sg*UXS1*) (Extended Data Fig. 5l). Knockout of *UGDH*, *PAPSS1,* or *UXS1* in lymphoma cells triggered a 48%, 48% or 60% depletion in DiI-LDL uptake relative to control cells, respectively. Altogether, our data shows that the synthesis of sulfated GAGs by cancer cells promotes ferroptosis resistance and an increase in the uptake of lipoproteins.

## Lipoprotein-mediated resistance to ferroptosis in tumours is driven by GAGs attached to cell surface proteoglycans

Sulfated GAGs are assembled in the Golgi apparatus, but they exert their functions in the extracellular space in two different modes: secreted forming part of the extracellular matrix, or localized at the cell surface^34^. One or more GAG chains are covalently linked to serine residues of a family of surface glycoproteins known as proteoglycans, through a tetrasaccharide linker^33^ (Fig. 3a). In the case of HS- and CS-proteoglycans, this tetrasaccharide is always initiated with addition of a xylose, followed by two galactose monosaccharides, and one GlcUA. At least 4 genes encoding for synthesis or glycosyltransferases enzymes involved in the formation of the linkage tetrasaccharide scored as essential for both ferroptosis resistance and lipoprotein uptake in our previous CRISPR screens (Fig. 2d, 3a): *UGDH*, *UXS1*, xylosyltransferase 2 (*XYLT2)*, and β-1,4-galactosyltransferase 7 (*B4GALT7*). This raises the possibility that cell surface GAGs covalently-bound to proteoglycans drive lipoprotein uptake and ferroptosis resistance in cancer cells.

**Figure 3.**
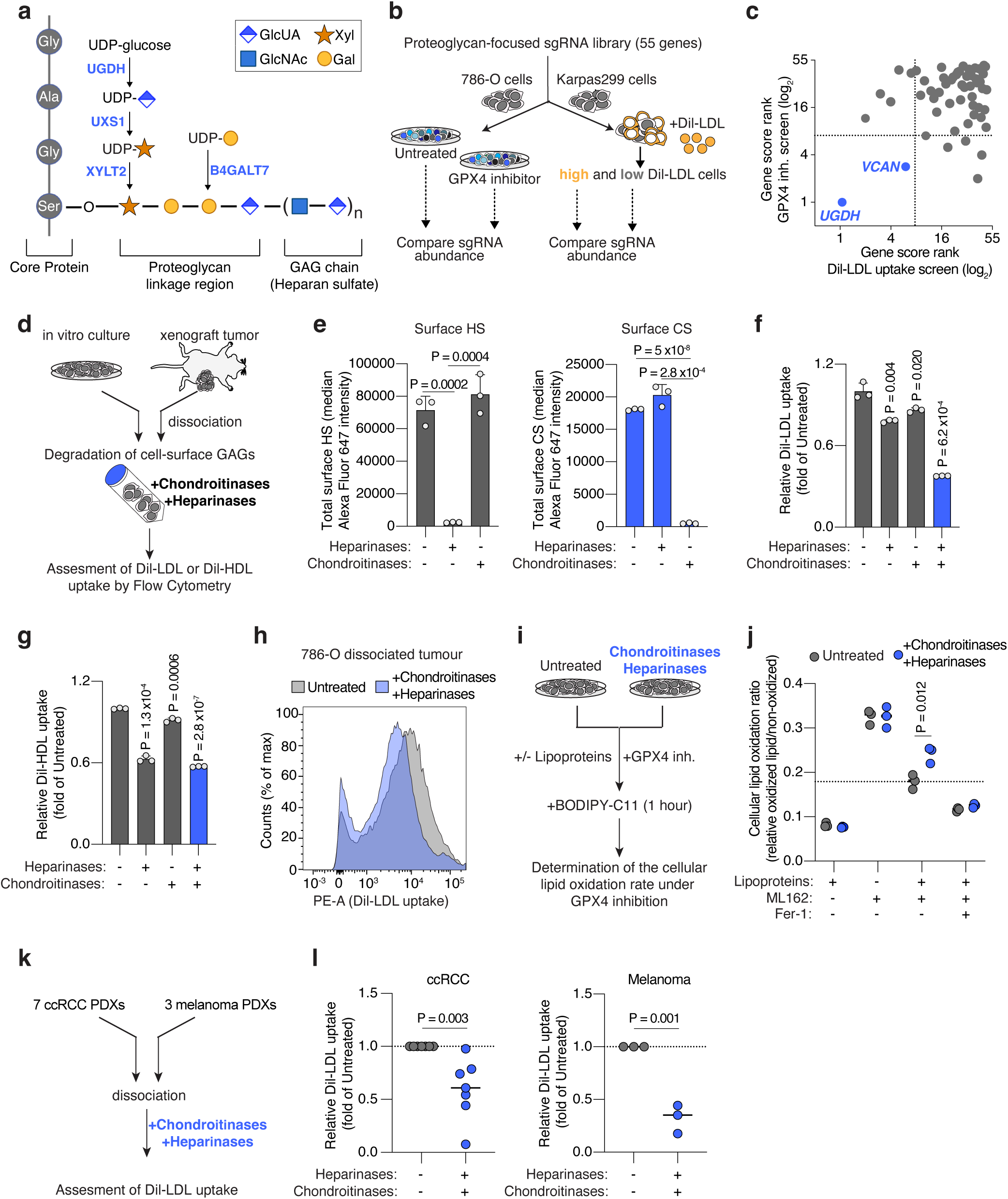
Cell-surface glycosaminoglycans bound to proteoglycans drive the uptake of LDL and HDL by cancer cells thus promoting ferroptosis resistance. **a.** Scheme of the canonical proteoglycan linkage region showing a heparan sulfate chain attached to a core protein through O-glycosylation of a serine residue. Essential genes for glycosaminoglycan-chain attachment to proteoglycans that scored in Karpas299 genetic screens are highlighted (blue). **b.** Schematic of CRISPR screens in 786-O (ccRCC) and Karpas299 (lymphoma) cells transduced with proteoglycan-focused sgRNA library (55 genes). 786-O cells were subjected to a proliferation-based screen in the presence or absence of the GPX4 inhibitor, ML162 (left). Karpas299 cells were incubated with DiI-LDL for 2 hours and subjected to FACS for high and low fluorescent cells. **c.** Plot of the gene score ranks in the 786-O and Karpas299 proteoglycan focused screens. Genes essential in both screens are highlighted (blue). sgRNAs targeting *UGDH* were included as a positive control. **d.** Experimental approach to degrade cell-surface HS and CS using heparinases and chondroitinases, respectively, on cancer cells in culture or dissociated from xenograft tumours. **e.** Total cell surface HS (left, grey) and CS (right, blue) in 786-O cells before and after treatment with heparinases (0.1 U/mL) and chondroitinases (0.1 U/mL) assessed by flow cytometry. **f.** Fold change in DiI-LDL uptake of 786-O cells upon treatment with heparinases (0.1 U/mL), chondroitinases (0.1 U/mL), or both (blue), relative to uptake of untreated cells, assessed by flow cytometry. **g.** Fold change in DiI-HDL uptake of 786-O cells upon treatment with heparinases (0.1 U/mL), chondroitinases (0.1 U/mL), or both (blue), relative to uptake of untreated cells, assessed by flow cytometry. **h.** Flow cytometry plot of the uptake of DiI-LDL in a cell suspension of 786-O xenograft tumours resected from mice left untreated (grey) or after combined treatment with heparinases and chondroitinases (blue). **i.** Experimental approach to evaluate ferroptosis sensitivity after enzymatic degradation of cell surface HS and CS. Cells treated with heparinases and chondroitinases are placed in media with or without lipoproteins and subjected to GPX4 inhibition, prior to staining with BODIPY-C11 and flow cytometry analysis of cellular lipid oxidation. **j.** Cellular lipid oxidation ratio using BODIPY-C11 of 786-O cells left untreated (grey) or after treatment with heparinases and chondroitinases (blue), and in the indicated conditions of lipoprotein availability and GPX4 inhibition (ML162, 200 nM). Fer-1 (1 μM) is included as a positive control of lipid peroxidation inhibition. **k.** Experimental approach to evaluate HS- and CS-mediated lipoprotein uptake in patient derived xenografts (PDXs). Fresh tissues were dissociated before treatment with or without heparinases and chondroitinases, and assessed for differences in DiI-LDL uptake using flow cytometry. **l.** Fold change in DiI-LDL uptake by ccRCC (left) and melanoma (right) PDX cell suspensions treated with heparinases and chondroitinases (1 U/mL of both, dark blue) relative to untreated samples (grey). Each dot represents a different PDX. **e-g,** Bars represent mean ± s.d.; **j, l,** bars represent the median in n=7 (ccRCC, left) and n=3 (melanoma, right) biologically independent samples; **e-g,** n=3 biological replicates. Statistical significance determined by two-tailed unpaired t-tests as indicated or compared to untreated cells (**f,g**).

Proteoglycans are a group of about 60 genes with diverse functions^22^. Whether specific HS- or CS-proteoglycans promote lipoprotein uptake and ferroptosis resistance in tumours, however, is unknown. Because sgRNAs against proteoglycan genes were not included in the metabolism-focused sgRNA library used in previous genetic screens, we constructed a proteoglycan-focused sgRNA library (10 sgRNAs per gene, 55 genes total). We transduced ccRCC (786-O) and lymphoma (Karpas299) cells with this proteoglycan library, and applied two complementary functional genetic strategies (Fig. 3b). 786-O cells were cultured for 14 population doublings in the presence or absence of a GPX4 inhibitor; whereas Karpas299 cells were incubated with DiI-LDL and sorted into top 5% and bottom 5% fluorescence populations. After analysis of sgRNA abundance in each condition, we defined gene essentiality (Extended Data Fig. 6a, 6b) and focused on genes that were essential in both screens (Fig. 3c). The proteoglycan-focused library included sgRNAs targeting *UGDH* to have a comparator for the effect of specific proteoglycans on ferroptosis resistance and LDL uptake. As expected, *UGDH* was the top hit in both genetic screens (Fig. 3c, Extended Data Fig. 6a, 6b). Among proteoglycan genes, the most essential in both screens was versican^35^ (*VCAN*) (Fig. 3c, Extended Data Fig. 6a, 6b), which encodes a large secreted proteoglycan decorated with multiple CS chains that can interact with hyaluronic acid^36^ and cell surface proteins.. Of note, the gene essentiality of *VCAN* for lipoprotein uptake and ferroptosis resistance in these CRISPR screens is significantly lower than *UGDH* essentiality.

To directly test the anti-ferroptotic role of VCAN in tumours, we next used CRISPR to disrupt *VCAN* expression in 786-O and Karpas299 cells (Extended Data Fig. 6c). Cells transduced with a sg*VCAN* lentivirus were more sensitive to GPX4 inhibition in proliferation experiments than sgControl isogenic cells (Extended Data Fig. 6d, 6e). Moreover, Karpas299 *VCAN*_KO cells were less efficient in taking up DiI-LDL (15% reduction, Extended Data Fig. 6f) and in growing as tumours in the flank of mice (Extended Data Fig. 6g) than cells expressing VCAN. Knocking out the CS proteoglycan *VCAN* in cancer cells, however, did not phenocopy the effects on lipoprotein uptake, ferroptosis resistance, or tumour growth observed in *UGDH*_KO cells. This suggests that other tumour proteoglycans promote the uptake of antioxidant-rich lipoproteins in *VCAN*_KO cells. Moreover, the genetic evidence for GAG biosynthesis regulating lipoprotein uptake and ferroptosis resistance suggests that both HS and CS could contribute to these phenotypes. We thus considered the possibility that, to impair the tumour uptake of antioxidant-rich lipoproteins, both HS and CS proteoglycans need to be targeted simultaneously.

To test the direct contribution of HS and CS to lipoprotein uptake, we designed an experimental approach to acutely deplete sulfated GAGs exclusively from the plasma membrane in wild-type cells. We dissociated ccRCC cells that were growing in culture with a non-enzymatic method that preserves membrane GAGs, and incubated these membrane-intact cancer cells with the following GAG-degrading enzymes^37^: bacterial heparinases (degrades HS), bacterial chondroitinases (degrades CS), or both (Fig. 3d). First, we employed HS or CS antibodies to label each cell surface GAG, respectively, and used flow cytometry to show that treatment of ccRCC cell lines 786-O and A-498 with heparinases or chondroitinases degraded surface HS or CS, respectively (Fig. 3e, Extended Data Fig. 7a, 7b). This effect was GAG-specific in that treatment with heparinases did not significantly change levels of surface CS and chondroitinases-treated cells had similar levels of HS as untreated cells (Fig. 3e, Extended Data Fig. 7a, 7b).

Next, we employed flow cytometry to assess DiI-LDL or DiI-HDL uptake capacity in cells with negligible levels of either surface HS or CS, or both, compared to control cells with intact HS and CS in their membranes. Degrading HS or CS decreased DiI-LDL uptake in 786-O cells by 22% and 14%, respectively (Fig. 3f), but the combined treatment with heparinases and chondroitinases reduced LDL uptake by 62% (Fig. 3f). Similar effects on DiI-LDL uptake were obtained using A-498 cells (Extended Data Fig. 7c). As for HDL uptake, we observed an additive effect of HS and CS-degradation in 786-O cells, with the combined treatment triggering a 43% reduction in cellular DiI-HDL relative to untreated cells (Fig. 3g). Surface HS and CS, however, did not impact DiI-HDL uptake of A-498 cells (Extended Data Fig. 7d), suggesting lipoprotein class-specificity and heterogeneity across cell lines.

The surface glycans of cancer cells may change between cell culture and the in vivo environment. To test whether the effect of surface GAGs on lipoprotein uptake is conserved in tumours, we injected 786-O cells in the flank of immunodeficient mice and let the tumours grow for two weeks. When subcutaneous tumours were palpable, we resected tumours and dissociated them into single cells using mechanical, non-enzymatic dissociation. We then performed DiI-LDL uptake assays in tumour-dissociated cells treated with or without heparinases/chondroitinases (Fig. 3d). Notably, degradation of HS and CS triggered a 50% decrease in the cellular internalization of DiI-LDL measured by flow cytometry (Fig. 3h, Extended Data Fig. 7e). Altogether, these experiments show that both HS and CS in the cell surface of tumours promote lipoprotein uptake, with their combined degradation triggering a decreased uptake similar to that observed upon inhibition of GAG biosynthesis by knocking out *UGDH*.

Lastly, we aimed to show that the increased lipoprotein uptake of cancer cells mediated by surface GAGs inhibits lipid peroxidation. We devised an assay in which 786-O cells were treated with both heparinases and chondroitinases, or left untreated (Fig. 3i). Cells with or without surface GAGs were then subjected to chemical GPX4 inhibition followed by addition of the lipid peroxidation fluorescent sensor BODIPY-C11. To test the contribution of lipoproteins, we performed this assay in the presence or absence of lipoproteins. Consistent with previous experiments showing the potent lipid antioxidant effect of lipoproteins, 786-O cells depleted of lipoproteins had a higher level of oxidized lipids upon GPX4 inhibition than cells with lipoproteins available (Fig. 3j). Heparinases/chondroitinases-treated cells, however, displayed higher lipid oxidation than untreated 786-O cells (Fig. 3j), but only when lipoproteins were present. These data suggests that both HS and CS on the surface of cancer cells promote resistance to ferroptosis through increased uptake of antioxidant-rich lipoproteins.

## Patient-derived xenografts employ surface GAGs to take up lipoproteins

We next investigated whether cell surface GAGs play a role in lipoprotein uptake in patient-derived xenografts (PDXs). We collected fresh tumours growing in mice from seven ccRCC and three melanoma PDXs, degraded surface GAGs, and assessed DiI-LDL uptake (Fig. 3k). Remarkably, ccRCC PDXs displayed a 40% decrease in LDL uptake upon combined treatment with heparinases and chondroitinases compared to untreated tumours (Fig. 3l). The LDL uptake capacity of melanoma PDXs depleted of HS and CS was reduced by 68% (Fig. 3l). Collectively, our data shows that GAGs are a major determinant of lipoprotein uptake in cancer cells in culture, tumour xenografts, and pre-clinical PDX models of ccRCC and melanoma.

## Human ccRCCs accumulate GAGs and lipoprotein-derived antioxidants

ccRCCs are characterized by prominent accumulation of cellular lipids^38^ and high rates of lipid uptake^4^. Moreover, we have shown that decreasing the levels of GAGs in three RCC patient-derived cell lines (A-498, Caki-2 and 786-O) sensitizes them to ferroptosis. To explore the role of GAGs on lipoprotein uptake in human kidney tumours, we collected paired specimens of ccRCC and adjacent kidney cortex tissues from 20 patients (Fig. 4a) and quantified levels of vitamin E, HS and CS.

**Figure 4.**
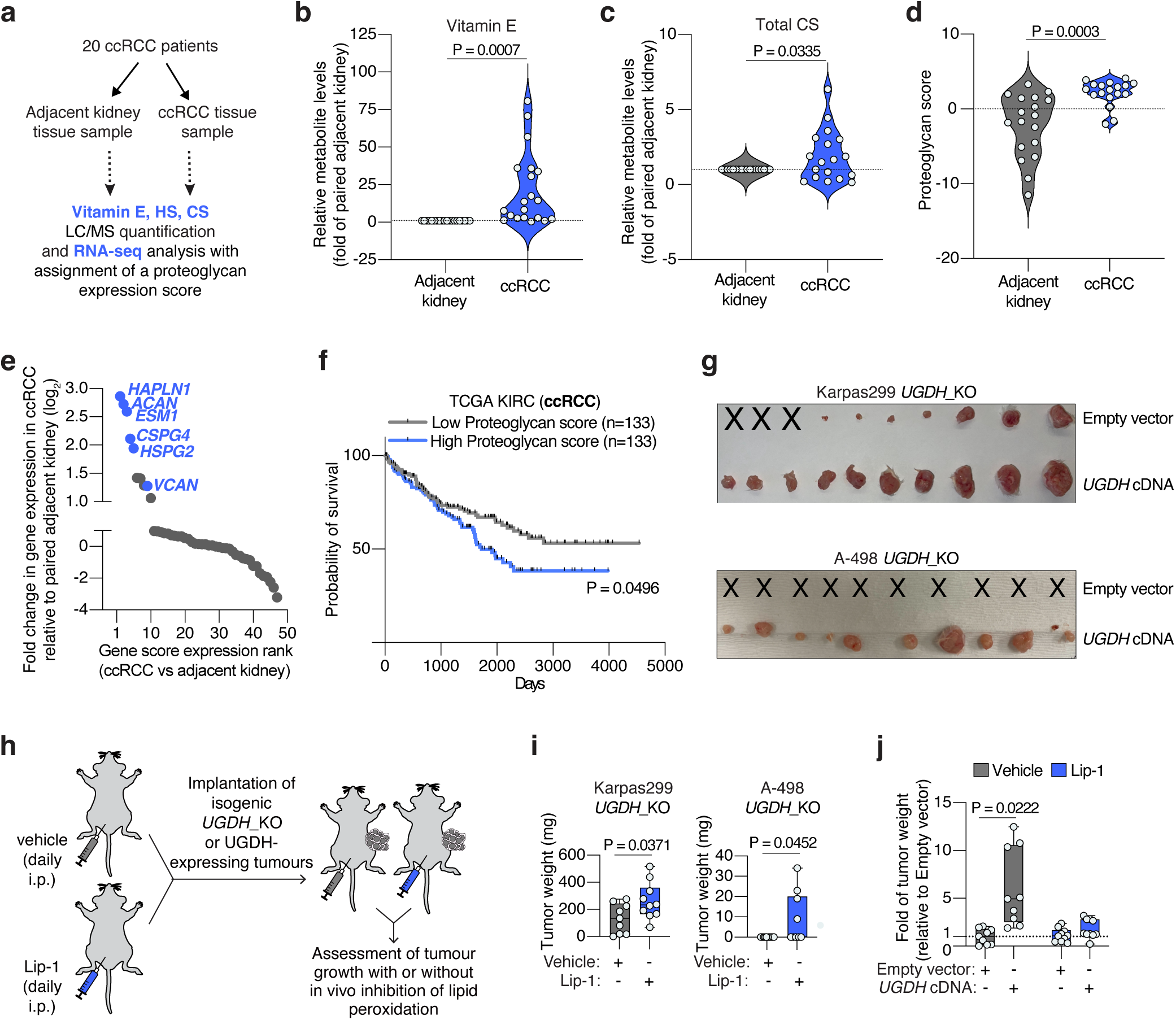
GAG-mediated lipoprotein uptake in pre-clinical models and human tissues. **a.** Primary ccRCC patient samples (adjacent kidney or tumour tissue) were subjected to parallel LC/MS quantification of vitamin E, HS or CS, and RNA-seq analysis. **b.** Violin plot showing the relative levels of vitamin E in ccRCC patient tissues (blue) compared to paired adjacent kidney (grey). **c.** Violin plot showing the relative levels of total CS per gram of protein in ccRCC patient tissues (blue) compared to paired adjacent kidney (grey). **d.** Violin plot showing the increased proteoglycan score of ccRCC patient tissues (blue) relative to paired adjacent kidney (grey) derived from RNA-seq analysis. **e.** Rank of proteoglycans by fold change in gene expression in ccRCC tumours relative to paired adjacent kidney. Relevant proteoglycans are highlighted in blue. **f.** Survival data in ccRCC patients (TCGA) stratified by high (top 25%, n=133, blue) or low (bottom 25%, n=133, grey) proteoglycan score. **g.** Representative tumour images of Karpas299 and A-498 *UGDH*_KO cells expressing a sgRNA resistant *UGDH* cDNA or an empty vector, and implanted subcutaneously in 6-12 week old immunodeficient mice. Injections that resulted in no engrafted tumour are marked “X”. **h.** Scheme showing daily intraperitoneal treatment of mice with vehicle (grey) or the anti-ferroptotic compound liproxstatin-1 (Lip-1, blue) before and after the implantation of *UGDH*_KO cells expressing a *UGDH* cDNA or an empty vector in mice. **i.** Tumour weights resulting from implantation of the indicated *UGDH*_KO cell lines on immunodeficient mice treated with vehicle (grey) or Lip-1 (blue) through daily intraperitoneal injection. **j.** Fold change in tumour weight formed by implantation of Karpas299 *UGDH*-KO cells expressing an *UGDH* cDNA or an empty vector in mice treated with vehicle (grey) or Lip-1 (blue) relative to vehicle-treated empty vector-transduced *UGDH*_KO tumours. **h, i,** Boxes represent the median, and the first and third quartiles, and the whiskers represent the minimum and maximum of all data points. **h, i**, n=10 biological replicates; **b-d**, n=18-20 biologically independent samples. Statistical significance determined by two-tailed unpaired t-tests.

Building upon the observation that most vitamin E is derived from lipoprotein uptake (Fig. 1h), we used this lipid as a proxy measurement of lipoprotein uptake in patient-derived tissues. Of the 20 pairs of specimens we analyzed, 17 had a higher than 2-fold increase in vitamin E in the tumour relative to the kidney, with ccRCCs containing a median 10.9-fold higher vitamin E levels than adjacent kidney (Fig. 4b). We also quantified free cholesterol in these samples as an internal lipid control, and found that cholesterol levels were only 1.7-fold higher in ccRCCs relative to adjacent kidneys (Extended Data Fig. 8a).

Next, we quantified the levels of HS and CS in paired samples and normalized values in ccRCCs to those in adjacent kidney. This analysis revealed an opposing pattern for each sulfated GAG. Total CS levels in cancerous tissues were higher than those in paired samples (Fig. 4c), whereas we found a decrease on total HS levels in ccRCCs (Extended Data Fig. 8b). Consistent with previous reports measuring GAGs on ccRCC patients^39^, our data suggests that human ccRCCs upregulate CS synthesis and decrease HS levels.

Collectively, our analysis shows that ccRCCs accumulate CS and vitamin E, suggesting that these tumours exacerbate synthesis of CS and lipoprotein uptake compared to the non-transformed kidney cortex. We next wanted to test whether these changes correlate with higher expression of proteoglycans, particularly CS-proteoglycans (CSPGs). We performed RNA sequencing in the same paired samples of ccRCC and adjacent kidney that we used for vitamin E and CS quantification (Fig. 4a). We focused our analysis on proteoglycans because of their more restricted expression than the ubiquitously expressed enzymes in the GAG biosynthesis pathway. A list of proteoglycan genes was subjected to a principal component analysis and the first principal component (PC1) was extracted and termed a ‘proteoglycan score’ (Extended Data Fig. 8c). ccRCC samples had a higher proteoglycan score than adjacent kidney samples (Fig. 4d). Moreover, this analysis revealed that several of the most upregulated proteoglycans in kidney tumours were potentially associated with higher lipoprotein uptake and ferroptosis resistance (Fig. 4e). Remarkably, among the five genes most upregulated in ccRCCs compared to adjacent kidney were three CSPGs: aggrecan (*ACAN*), endocan (*ESM1*), and *CSPG4*. The other two genes in this group were Hyaluronan And Proteoglycan Link Protein 1 (*HAPLN1*), which interacts with VCAN^40^; and perlecan (*HSPG2*), a large HS- and CS-bound proteoglycan that contains a LDLR-like domain in its structure and interacts with LDL^41^. VCAN was also upregulated in tumour tissues (Fig. 4e, Extended Data Fig. 8c).

Lastly, we analyzed the TCGA ccRCC dataset (KIRC) for proteoglycan scores and compared patient survival of patients with a high (top 25%) and low (bottom 25) proteoglycan score. Patients with a high proteoglycan score had poor survival compared to the low proteoglycan score group (Fig. 4f). Therefore, the mass spectrometry and gene expression analyses of ccRCC patient specimens compared to the adjacent kidney showcases that CS levels and RNA levels of associated proteoglycans strongly correlate with high accumulation of lipoprotein-derived antioxidants and poor patient survival.

## GAGs promote tumour growth through their anti-ferroptotic effect

Oxidative stress, including lipid oxidative stress, can be a metabolic limitation for tumour growth and progression^3,9,11,42^. We thus tested whether loss of tumour GAGs impacts cancer growth in mice. *UGDH*_KO Karpas299, A-498 and HeLa cells injected as subcutaneous tumours formed significantly smaller tumours than *UGDH*-expressing isogenic controls (Fig. 4g, Extended Data Fig. 9a, 9b). Karpas299 *UGDH*_KO and HeLa *UGDH*_KO tumours were 3.9-fold and 2.1-fold smaller than isogenic tumours expressing a *UGDH* cDNA, whereas A-498 *UGDH*_KO tumours did not engraft but formed palpable tumours when expressing UGDH.

To confirm that the loss of GAGs impairs growth through increased lipid peroxidation, we treated mice daily with a lipophilic antioxidant (liproxstatin-1, Lip-1)^43^ before and after the implantation of *UGDH*_KO or UGDH-expressing isogenic tumours (Fig. 4h). Treatment with Lip-1 stimulated growth of *UGDH*_KO Karpas299 and A-498 tumours, but not tumours that express UGDH (Fig. 4i, Extended Data Fig. 9c). Supplementation with Lip-1 allowed A-498 *UGDH*_KO tumours to develop, though it did not fully restore their growth to levels seen in UGDH-expressing counterparts (Fig. 4j, Extended Data Fig. 9c). Taken together, our data shows that GAG biosynthesis plays a critical role in sustaining tumour growth by promoting lipoprotein-dependent resistance to lipid peroxidation.

## DISCUSSION

The anti-ferroptotic role of some antioxidant lipids has previously been reported, but their relevance to in vivo biology is unknown. These lipids are studied in the context of either de novo synthesis or individual lipid supplementation in culture at supraphysiological levels. The crucial transport system for these lipids, lipoproteins, is largely ignored in cell culture studies where lipoproteins in fetal bovine serum (FBS) are well below physiologically relevant lipoprotein concentrations. We discover that lipoproteins supplemented in cell culture are a major antioxidant reservoir for cancer cells. Whether dysregulated levels of circulating lipids in humans affect the tumour antioxidant response and cancer progression needs further testing.

We unexpectedly find that lipoprotein uptake in cancer cells depends on cell-surface sulfated GAGs. GAGs promote the uptake of at least two classes of lipoproteins, LDL and HDL, through their binding to proteoglycans. Recent evidence suggests that certain proteoglycans in hepatocytes and macrophages can promote LDL uptake^44,45^, which raises the possibility that the GAG-mediated mechanism described herein may be more widely applicable to other physiological systems and disease context involving dysregulated lipid metabolism.

The CRISPR screen on lymphoma cells revealed the essentiality of two genes essential for GAG sulfation in the uptake of antioxidant-rich lipoproteins. This is consistent with previous reports of the role of GAG sulfation in macrophages during atherosclerosis^46^. Building upon this, the focus of our experiments were the two major classes of sulfated GAGs, HS and CS. We have not formally tested the relevance of another major, non-sulfated GAG on lipoprotein uptake: hyaluronic acid (HA). HA synthesis also depends on the enzymatic product of UGDH, GlcUA. Although HA synthesis enzymes did not score in our initial genetic screen, VCAN interacts with HA and the HA-proteoglycan HAPLN1, whose RNA levels are upregulated in human ccRCCs compared to adjacent kidney. It is possible that HA plays a supporting role in the CS- and HS-mediated effect on lipoprotein uptake by cancer cells.

In patients with ccRCC, high proteoglycan gene expression correlates with worse survival, suggesting that GAGs and proteoglycans could contribute to cell fitness by promoting antioxidant-rich lipoprotein uptake. Lipoproteins may be a readily accessible source of crucial antioxidants that enable the survival of cancer cells under metabolic stress. Hence, targeting GAGs in tumours could be a powerful strategy to deplete cancer cells of key antioxidants and promote tumour ferroptosis even in conditions of high lipoprotein availability and diets with high fat contents.

## MATERIAL AND METHODS

### Compounds

The following compounds were used: ferrostatin-1, liproxstatin-1, ML162, oleic acid (Cayman Chemical); puromycin, blasticidin, BODIPY-C11 581/591, cholecalciferol (vitamin D_3_), protease inhibitor cocktail (Fisher Scientific); RPMI-1640, Hanks’ Balanced Salt Solution (HBSS) (Gibco); purified human low density lipoproteins (LDL), purified human high density lipoproteins (HDL), purified human DiI-LDL, purified human DiI-HDL, lipoprotein-depleted fetal bovine serum (LPDS) (Kalen Biomedical); Bacteroides Heparanase I, Bacteroides Heparanase II, Bacteroides Heparanase III, T4 Ligase, BamHI, NotI, BsmBI (New England Biolabs); Cultrex BME, Type 3 (R&D Systems); coenzyme Q10 (CoQ), ML210, erastin (Selleck Chemical); bovine serum albumin (BSA), chondroitinase ABC from proteus vulgaris, cholesterol, alpha-tocopherol (vitamin E), menaquinone-4 (vitamin K_2_), ammonium iron (III) citrate (FAC), dimethyl sulfoxide (DMSO), tween-80, polybrene, polyethylene glycol-300 (PEG-300), fetal bovine serum (FBS), non-enzymatic cell dissociation solution, DAPI (Sigma-Aldrich); TransIT-LT1 transfection reagent (MirusBio).

Antibodies to ACSL3 (Ab151959), human GPX4 (Ab41787), LDLR (52818), SCARB1(52629) were purchased from Abcam; Heparan Sulfate (370255-S) from AMSBio; IRP2 (37135S), SLC7A11 (12691S), TFRC (13208S), and β-tubulin (2146S) from Cell Signaling Technology; GAPDH (GTX627408) and UGDH (GTX104993) from GeneTex; AIFM2 (20866-1-AP) and PAPSS1 (14708-1-AP) from ProteinTech; mouse GPX4 (sc-166570) from Santa Cruz Biotechnology; and Chondroitin Sulfate (C8035) from Sigma. Horseradish peroxidase (HRP)-conjugated anti-rabbit (7074S) and anti-mouse (7076S) antibodies were purchased from Cell Signaling Technology and goat anti-mouse Alexa Fluor 647 (A-21238) antibody was purchased from Invitrogen.

### Cell lines and cell culture

All cell lines were purchased from ATCC, authenticated by short tandem repeat profiling, and verified to be mycoplasma-free every 2 months. They were maintained at 37°C and 5% CO_2_ and cultured in RPMI-1640 medium supplemented with 1mM glutamine, 10% fetal bovine serum, penicillin and streptomycin. For experiments performed in the absence of serum lipoproteins, RPMI-1640 medium was supplemented with 10% lipoprotein-depleted fetal bovine serum (Kalen Biomedical) in place of the normal 10% fetal bovine serum. For LDL or HDL supplementation experiments, we used the indicated concentrations of purified human LDL or HDL (Kalen Biomedical).

### Constructs

For generation of lentiviral knockouts constructs, annealed oligonucleotides (below) were cloned into lentiCRISPR-v2 vector using T4 Ligase.

sg*ACSL3*_2 forward, 5’-CACCGGGGCTGGAACAATTTCCGA-3’;

sg*ACSL3*_2 reverse, 5’-AAACTCGGAAATTGTTCCAGCCCC-3’;

sg*AIFM2*_5 forward, 5’-CACCGGATAAGATGAGAGAAGGGC-3’;

sg*AIFM2*_5 reverse, 5’-AAACGCCCTTCTCTCATCTTATCC-3’;

sg*GPX4*_1 forward, 5’-CACCGTAGGCGGCAAAGGCGGCCG-3’;

sg*GPX4*_1 reverse, 5’-AAACCGGCCGCCTTTGCCGCCTAC-3’;

sg*Gpx4*_1 forward, 5’--3’; CACCGCGTGTGCATCGTCACCAACG

sg*Gpx4*_1reverse, 5’--3’; AAACCGTTGGTGACGATGCACACG

sg*LDLR*_5 forward, 5’-CACCGTGGCCCAGCGAAGATGCGA-3’;

sg*LDLR*_5 reverse, 5’-AAACTCGCATCTTCGCTGGGCCAC-3’;

sg*PAPSS1*_2 forward, 5’-CACCGAACAAGAGAGGTCAGGTGG-3’;

sg*PAPSS1*_2 reverse, 5’-AAACCCACCTGACCTCTCTTGTTC-3’;

sg*SCARB1*_2 forward, 5’-CACCGCTGGAGTTCTACAGCCCGG-3’;

sg*SCARB1*_2 reverse, 5’-AAACCCGGGCTGTAGAACTCCAGC-3’;

sg*UGDH*_7 forward, 5’-CACCGCTCTGCCAGAAACTCAGGGT-3’;

sg*UGDH*_7 reverse, 5’-AAACACCCTGAGTTTCTGGCAGAGC-3’;

sg*Ugdh*_7 forward, 5’- -3’; CACCGAAGTAGTCGAATCCTGTCG

sg*Ugdh*_7 reverse, 5’- -3’; AAACCGACAGGATTCGACTACTTC

sg*UXS1*_8 forward, 5’-CACCGAGGTCCTATTGGATTCACG-3’;

sg*UXS1*_8 reverse, 5’-AAACCGTGAATCCAATAGGACCTC-3’;

sg*VCAN*_4 forward, 5’-CACCGCAGTAAATTCACCTTCGAGG-3’;

sg*VCAN*_4 reverse, 5’-AAACCCTCGAAGGTGAATTTACTGC-3’

Guide resistant *UGDH* cDNA was synthesized as gene fragments (TWIST Biosciences). PCR-overlap extension and Gibson assembly were used to clone the cDNA into PMXS-IRES-Blast.

### Generation of knockout and cDNA overexpression cell lines

For virus production, sgRNA or cDNA-containing vector was transfected into HEK293T cells with packaging vectors for lentivirus (VSV-G and Delta-VPR) and retrovirus (VSV-G and Gag-pol), respectively, using TransIT-LT1 Transfection Reagent. 24 hours after transfection, the transfection medium was replaced with fresh medium. 48 hours after transfection, medium containing viral particles was collected and filtered using a 0.45-mm filter to exclude HEK293T cells. 24 hours before infection, cells of interest were plated in six-well tissue culture plates (200,000 cells per well for suspension lines and 100,000 cells per well for adherent lines). For transduction, infection medium and 8 µg/ml of polybrene were added and cells were spin infected at 2,200rpm for 1.5 hours. 24 hours after infection, infection medium was replaced with fresh medium. To initiate selection of transduced cells, puromycin (for sgRNA lentiviral vector) or blasticidin (for overexpression retroviral vectors) was added 24 hours after addition of fresh medium. For *UGDH/Ugdh*_KO cells and *GPX4*/*Gpx4*_KO cells, sgRNAs targeting *UGDH/Ugdh* or *GPX4*/*Gpx4* were cloned into lentiCRISPR-v2. After transduction and selection using puromycin, single cell clones were plated in medium containing 1 µM ferrostatin-1. Knockout and/or overexpression of the target gene was verified by immunoblot analysis. Cells were then maintained in media without ferrostatin-1.

### Immunoblotting

Cell pellets were washed twice with PBS and lysed in lysis buffer (10mM Tris-Cl pH 7.5, 150 mM NaCl, 1mM EDTA, 1% Triton X-100, 2% SDS, and 0.1% CHAPS) supplemented with protease inhibitors. After a sonication and a 10-min incubation on ice, cell lysates were centrifuged for 10 min at 13,000g and 4°C. Supernatants were collected from each sample, and protein concentration was determined via a Pierce BCA Protein Assay Kit (Thermo Scientific). BSA was used as a protein standard. Lysates were diluted to a total of 20 µg of protein in 20 µL of lysis buffer and resolved on 4-12, 10–20, or 12% SDS– PAGE gels before transfer to Immobilin-P polyvinyl difluoride membrane (Millipore). Membranes were blocked in 5% non-fat milk in TBS-T and analyzed using standard immunoblotting protocols.

### Proliferation assays

Cell lines were cultured in triplicates in 96-well plates at 1,000 (suspension cell lines) or 500 (adherent cell lines) cells per mL in a final volume of 0.2 mL medium with the indicated treatments. Untreated cells were plated and read at the initial timepoint for use in data normalization. After five days of growth, 40 µL of CellTiter-Glo reagent (Promega) was added to each well and luminescence was read using an Infinite M Plex plate reader (Tecan). Data are presented as the relative fold change (log_2_) in luminescence to that of the initial number of cells.

### CRISPR-based genetic screens

The highly focused metabolism^47^ and the metabolism human^19^ sgRNA libraries have been generated and used before. The proteoglycan-focused sgRNA library was designed as previously described by curating a list representation of bona-fide identified proteoglycan genes. Oligonucleotides (IDT) were annealed before cloning into lentiCRISPR-v2 using a T4 Ligase. This plasmid pool was used to generate a lentiviral library, which was transfected into HEK293T cells and used to generate viral supernatant as described above. Cells were infected at a multiplicity of infection of 0.7 and selected with puromycin. An initial sample of 3 million (focused-metabolism and proteoglycan-focused sgRNA library) or 30 million cells (metabolism sgRNA library) were harvested and infected cells were cultured for 14 population doublings under the indicated conditions (RPMI containing LPDS and 5 µg/mL cholesterol; RPMI containing LPDS, 5 µg/mL cholesterol, and 20 µg/mL HDL, 20 µg/mL LDL; sublethal ML210, 0.5 *µ*M; sublethal ML162, 0.5 µM). Final samples of 3 or 30 million cells were collected and genomic DNA was extracted (DNeasy blood and Tissue kit, Qiagen). Next, sgRNA inserts were PCR-amplified using specific primers for each condition. PCR amplicons were then purified and sequenced on a NextSeq 2000 (Illumina). Sequencing reads were mapped and the abundance of each sgRNA was measured. The gene score for each gene was defined as the median log_2_ fold change in the abundance each sgRNA targeting that gene between conditions. A gene score lower than -1 is considered significant.

For fluorescence-activated cell sorting (FACS) DiI-LDL screens, cells were transduced, selected, and expanded following the above workflow. An initial pool of 3 or 30 million cells was collected. Transduced cells were placed in RPMI-medium with LPDS overnight the day before cell sorting. 2 hours prior to sorting, medium was replaced with HBSS containing 10 µg/ml of DiI-LDL. After a 2 hour DiI-LDL incubation, cells were collected, washed twice with HBSS, and resuspended in HBSS containing DAPI. The cell suspension was strained through a 40-um cell strainer (Falcon) and sorted on a FACSAria III (BD Biosciences). The top and bottom 5% of DiI-LDL fluorescing cells were collected via sorting. Sorted cells were maintained RPMI medium supplemented with ferrostatin-1 (1 µM), until a final pool of 3 or 30 million cells could be collected for analysis of sgRNA abundance.

### Flow cytometry determination of cellular DiI-LDL and DiI-HDL uptake

For cell culture experiments, cell lines were plated at a density of either 150,000 (adherent) or 300,000 (suspension) cells per well in six-well plates in RPMI medium with LPDS. After 24 hours, the medium was removed. For experiments involving heparinases and chondroitinases treatment, each well was washed with HBSS once prior addition HBSS containing 0.1 U/mL Heparinases I, II, III and 0.1 U/mL Chondroitinases for 2 hours at 37C. The medium/HBSS was then replaced with HBSS containing 10 µg/ml of DiI-LDL or DiI-HDL. Following a 2-4 hour incubation with the DiI-labeled lipoprotein, cells were collected, washed twice with cold HBSS, and resuspended in HBSS containing DAPI. The cell suspension was strained through a 40-μm cell strainer (Falcon) and maintained on ice until analysis.

Data acquisition was performed using a FACS Canto RUO (BD Biosciences). Data were collected from the phycoerythrin (PE) detector for DiI-LDL/DiI-HDL uptake and the shift in median fluorescence intensity of PE was analyzed. At least 10,000 events were recorded per sample. Data analysis was conducted using FlowJo v.10 software, gating initially for singlet events based on forward and side scatter characteristics and subsequently for live cells based on DAPI staining.

For ex vivo experiments, 786-O mouse xenograft tumours or fresh ccRCC and melanoma patient-derived xenograft tissue were dissociated using non-enzymatic cell dissociation solution (Sigma) under continuous rotation at 150rpm and 37°C for 2 hours in MACS Gentle Dissociation C-Tubes (Miltenyi Biotec) to form a cell suspension. This cell suspension was filtered using a 70 µM cell strainer (Falcon) and treated with 1X RBC Lysis Buffer (Invitrogen) at room temperature for 10 minutes to eliminate red blood cells. Tumour cells were then washed twice with HBSS. Cells were plated into 24 well tissue culture plates and left untreated (in HBSS alone) or treated with HBSS containing 1.0 U/mL Heparinases I, II, III and 1.0 U/mL Chondroitinases for 2 hours at 37°C and under continuous rotation (100 rpm). After glycosidase treatment, 20 µg/mL DiI-LDL was added to the suspension and cells were then incubated overnight at 37°C under continuous rotation (100 rpm). Following overnight incubation, cells were pelleted, washed twice with HBSS, and resuspended in HBSS containing DAPI. The cell suspension was strained through a 40-μm cell strainer (Falcon) and maintained on ice until analysis. Data was acquired and analyzed as above.

### Determination of cell-surface HS and CS Content

Cells were plated at 150,000 cells per well in six-well tissue culture plates in RPMI medium supplemented with LPDS 24 hours prior to analysis. After 24 hours, the medium was removed, each well was washed with HBSS, and the medium was replaced with HBSS containing 0.1 U/mL Heparinases I, II, III and 0.1 U/mL Chondroitinases. Cells were then incubated for 2 hours at 37°C. Following incubation, cells were dissociated using non-enzymatic cell dissociation solution (Sigma), collected, and washed with HBSS before incubation with antibodies against heparan sulfate (HS, AMSBio; 1:2000) and chondroitin sulfate (CS, Sigma; 1:2000) for 30 minutes on ice. Cells were then washed with HBSS before incubation with a secondary antibody anti-goat Alexa Fluor 647 (Invitrogen; 1:1000) for an additional 30 minutes on ice. Next, cells were washed twice with HBSS, resuspended in HBSS containing DAPI, passed through a 40-μm cell strainer (Falcon), and maintained on ice until analysis. Data acquisition was performed using a FACS Canto RUO (BD Biosciences). Data were collected from the Alexa Fluor 647 detector and the shift in median fluorescence intensity of Alexa Fluor 647 was analyzed. At least 10,000 events were recorded per sample. Data analysis was conducted using FlowJo v.10 software, gating initially for singlet events based on forward and side scatter characteristics and subsequently for live cells based on DAPI staining.

### Determination of cellular lipid peroxidation using BODIPY-C11

48 hours before collection, cells were plated at a density of 200,000 (adherent) or 300,000 (suspension) cells per well in six-well tissue culture plates in RPMI medium containing LPDS. 24 hours later the indicated cells were pre-treated by supplementation of Vitamin E (10 µM), Vitamin K2 (10 µM), Vitamin D3 (10 µM), Vitamin A (10 µM), CoQ10 (10 µM), Fer-1 (1uM), LDL (50 µg/ml), HDL (50 µg/ml) for 2 hours. After 2 hours, indicated cells were treated with the corresponding dose of ML162 to induce lipid peroxidation. After 24 hours of ML162 treatment, cells were collected, washed with HBSS, and incubated with BODIPY-C11 (2 µM) for 1 hour at 37°C in the dark. Cells were then washed twice with HBSS, resuspended in HBSS containing DAPI, passed through a 40-μm cell strainer (Falcon), and maintained on ice until analysis. Data acquisition was performed using a FACS Canto RUO (BD Biosciences). Data was collected from the FITC detector for oxidized BODIPY and the PE detector for reduced BODIPY. Data analysis was conducted using FlowJo v.10 software, gating initially for singlet events based on forward and side scatter characteristics and subsequently for live cells based on DAPI staining. The cellular lipid oxidation ratio was calculated as the ratio of oxidized BODIPY (FITC signal) to total BODIPY (FITC signal + PE signal).

For experiments in Figure 3i-j: Cells were plated at a density of 200,000 cells per well in six-well tissue culture plates in RPMI medium containing LPDS 24 hours prior to treatment/analysis. After 24 hours, the medium was removed from each well and replaced HBSS containing 0.1 U/mL Heparinases I, II, III and 0.1 U/mL Chondroitinases before incubation at 37°C for 2 hours. Following glycosidase incubation, each well was washed with HBSS and replaced with RPMI medium containing either LPDS or FBS. Cells were incubated in these media with different lipoprotein levels for 2 hours. After 2 hours, ML162 (200nM) was added to the indicated wells and cells were incubated for 2 hours. After 2 hours of ML162 incubation, cells were collected, washed with HBSS, and incubated with BODIPY-C11 (2 µM) for 1 hour at 37°C in the dark. Cells were then washed twice with HBSS, resuspended in HBSS containing DAPI, passed through a 40-μm cell strainer (Falcon), and maintained on ice until analysis. Data was acquired and analyzed as above.

### Subcutaneous xenograft tumours in mice

All animal studies were approved by the respective Institutional Animal Care and Use Committees (IACUC) at the University of Texas Southwestern Medical Center and animal care was conducted in accordance with institutional guidelines. All mice were maintained on a standard light-dark cycle with food and water ad libitum. The maximum tumour size allowed is 2 cm at the largest diameter or 10% of the animal’s body weight; this maximum size was not exceeded.

Xenograft tumours were initiated by injecting the following number of cells in DMEM with 30% Cultrex:

- 75,000 cells per 100 μL for Karpas299 parental and individual knockout cell lines.
- 75,000 cells per 100 μL for HeLa parental and individual knockout cell lines.
- 500,000 cells per 100 μL for A-498 parental and individual knockout cell lines.

After subcutaneous injection into the left and right flanks of male and female 6–14-week-old NOD SCID gamma (NSG) mice, tumours were grown for 2–8 weeks.

Additionally, we performed tumour growth experiments using the anti-ferroptotic compound liproxstatin-1 (Lip-1). Lip-1 was reconstituted in DMSO, followed by consecutive additions of PEG-300, Tween-80, and water to a final solvent mixture of 5.1% DMSO, 40% PEG-300, 2% Tween-80, 55.9% Water. The resulting solution was thoroughly vortexed and sonicated in a water bath until clear, followed by a centrifugation and storage as aliquots at -70°C. A solution with DMSO not containing Lip-1 was used as a control for the experiment. Vehicle or Lip-1 (10 mg/kg) solutions were administered through intraperitoneal injections from 3 days before tumour implantation and daily until the end of the experiment.

### LC/MS determination of vitamin E and cholesterol

For extraction of vitamin E, we used a non-polar extraction protocol previously established^48^ to maximize vitamin E extraction. Briefly, we collected 3×10^6^ HeLa cells or 50-100 mg of human ccRCC or adjacent kidney tissues, resuspended them in 800 µl of PBS, lysed using a BeadBlaster 24R (Benchmark Scientific) followed by sample sonication for 60 seconds. 1/10 of this solution was collected for protein quantification for normalization of values. The remaining supernatant was processed as following: addition of 700 µL of LC/MS grade EtOH + 2.1 mL of LC/MS grade hexane (Sigma). Solutions were then thoroughly vortexed for 5 minutes at 4°C. After centrifugation, the upper layer was collected into a new tube. Next, we re-extracted the remaining aqueous phase by adding 300 µL of EtOH + 900 µL of hexane, followed by vortexing and centrifugation. The two non-polar phases containing vitamin E and cholesterol were then collected together, dried down and stored at -70°C until analysis.

Before analysis, samples were stabilized at room temperature, followed by addition of HPLC 100% ethanol. Vitamin E and cholesterol were analyzed with a SCIEX QTRAP 5500 liquid chromatograph/triple quadrupole mass spectrometer coupled with a Nexera Ultra-High-Performance Liquid Chromatograph system (Shimadzu Corporation). Separation was achieved as previously described^49^ using a Phenomenex Synergi™ Polar RP HPLC column (150 × 2.0 mm, 4µm, 80 Å). Mobile phase A composition was 2.0 mM ammonium acetate in methanol/water (65/35, v/v) and mobile phase B composition was 2.0 mM ammonium acetate in methanol/isopropanol (63/37, v/v). The gradient elution was: 0–2.0 min, linear gradient 0–60% B, 2.0–5.6 min, linear gradient 60–100% B, then the column was washed with 100% B for 4.4 min before reconditioning it for 5 min using 0% B. The flow rate was 0.5 ml/min at 40°C, and the injection volume was 10 µL. The mass spectrometer was used with an atmospheric-pressure chemical ionization (APCI) source in multiple reaction monitoring (MRM) mode. We analyzed MRM data with Analyst 1.6.3 software (SCIEX). The MRMs (Q1/Q3) used for metabolites were 369.0/161.0 (cholesterol, CE: 15), 431.4/165.1 (Vitamin E, CE: 30), and 437.4/171.1 (Vitamin E-d_6_, internal standard, CE: 30).

### LC/MS determination of HS and CS

For purification and analysis of glycosaminoglycans, we adapted a previously established protocol^50^. Briefly, 5×10^6^ cells/genotype or 25-100 mg of human ccRCC or adjacent kidney tissues were collected for analysis. Cell pellets were solubilized by adding 0.5 mL of 1X RIPA buffer (Millipore). Lyophilized tissues were homogenized in 5 mL of ice-cold wash buffer containing 50 mM sodium acetate, 0.2 M NaCl, pH 6.0. A portion of each cell/tissue sample was diluted 1:10 in wash buffer and digested overnight at 37°C with 0.5 mg/mL Pronase (Sigma) and 0.1% Triton X-100 prior to HS/CS purification. GAGs were purified from cell/tissue homogenates using DEAE-Sepharose anion exchange chromatography, as described previously^51^. All preparations were desalted by gel filtration (PD-10, Cytiva). For HS disaccharide analysis, lyophilized GAGs were incubated with 2 mU each of heparin lyases I, II, and III (IBEX) for 16 h at 37°C in a buffer containing 40 mM ammonium acetate and 3.3 mM calcium acetate, pH 7. For CS disaccharide analysis, lyophilized GAGs were incubated with 2 mU of chondroitinase ABC (Sigma) for 16 h at 37°C in a buffer containing 500 mM Tris and 500 mM NaCl, pH 7.9. HS/CS disaccharides were aniline-tagged and analyzed by HILIC-LC-MS on a Waters SYNAPT XS mass spectrometer equipped with an ACQUITY UHPLC H-class system (BEH Glycan Column, 2.1 mm X 100 mm). Mobile phase A was 50mM ammonium formate buffer, pH 4.4. Mobile phase B was 100% acetonitrile. The elution was: 0–5.0 min with isocratic 10% A, linear gradient 10–33% A for 5.0–49.0 min, linear gradient 33–45% A from 49.0-51.5 min, isocratic 45% A from 52.0-60 min, then the column was washed with 90% B for 10 min. The column temperature was room temperature, and the flow rate used was 0.5 mL/min (injection volume of 2 µL). Total GAGs were normalized to protein amount measured via BCA assay (cells) or tissue weight.

### Determination of *UXS1* and *VCAN* knockout efficiency

Cells transduced with a sgControl or a sgRNA targeting *UXS1* or *VCAN* were collected and their genomic DNA extracted using QuickExtract DNA (Lucigen) following manufacturer’s instructions. A fragment of *UXS1* or *VCAN* genes was PCR-amplified using GoTaq DNA Polymerase (Promega) with the corresponding primers (see below). The PCR product was run on an agarose gel and purified by gel DNA recovery kit (Zymo Research), followed by cloning into pGEM-T vector by pGEM-T easy vector system (Promega). The ligation product was transformed into DH5α competent cells. We then added 20 μL of 500 mM IPTG solution and 20 μL of 50mg/ml X-Gal solution on LB plate containing 50 μg/mL carbenicillin disodium. After overnight incubation at 37°C, we picked white colonies for amplification and plasmid isolation by QIAprep Spin miniprep kit (QIAGEN). Lastly, we subjected the resulting vectors to Sanger sequencing followed by sequence alignment between isogenic cells to verify gene editing efficiency.

The following primers were used:

*UXS1* exon 9 forward, 5’-GGGTCTTTCCGAAGTTCCCT-3’;

*UXS1* exon9 reverse, 5’-GCCTAAACCGCAAGCCTAGA-3’;

*VCAN* exon 6 forward, 5’-ATCAGAAGCTGCTACCTTGCC-3’;

*VCAN* exon 6 reverse, 5’-CCCAAGTAGACCACCTCCAC-3’;

### RNA extraction and RNA-sequencing of kidney and ccRCC samples

RNA extraction was performed using the Trizol reagent (Thermo Fisher Scientific) followed by purification with the RNeasy Mini Kit (Qiagen). The concentration of total RNA was determined using the Qubit fluorometer and the Invitrogen Qubit RNA High Sensitivity kit (Invitrogen).

For RNA-seq library preparation, the NEBNext Ultra II directional RNA library prep kit with the NEBNext Poly(A) mRNA magnetic isolation module (New England Biolabs) was utilized according to the manufacturer’s protocol. Libraries were indexed using standard N.E.B indices (New England Biolabs). Sequencing reads were aligned to the human reference genome (hg19) using STAR 2.7.3.a in the 2-pass mod, and gene counts were generated with htseq-count v0.6.1. Differential expression analysis was conducted using DESeq2 v1.14.1.

Additionally, sequences were trimmed to remove adapters and sequences with quality scores <25. Reads shorter than 35bp after trimming were discarded. Alignment to the GRCh38 genome assembly was performed using HiSAT2, and duplicate reads were marked with SAMBAMBA. Feature counting (genes, transcripts, and exons) was carried out using featureCounts. Differential expression analysis was also performed using EdgeR and DESeq.

### Proteoglycan score calculations

A list of proteoglycan genes was manually curated. Principal component analysis was performed on log_2_-transformed, mean-centered, and z-transformed data, and the first principal component (PC1) was extracted and used as the Proteoglycan score, as previously described^52^. The same analysis was applied for the Cancer Genome Atlas Kidney Renal Clear Cell Carcinoma (TCGA-KIRC) dataset and patient samples were ranked by proteoglycan score.

### Patient-derived xenografts

All tumour specimens for patient derived xenografts were obtained after patients signed informed consent documents for studies approved by the Institutional Review Board (IRB) at the University of Texas Southwestern Medical Center (UTSW) or the University of Michigan.

Patients diagnosed with renal cell carcinoma (RCC) were consented in accordance with our approved Institutional Review Board study protocol (STU012011-190). Patient-derived xenografts studies were conducted according to the UT Southwestern Institutional Animal Care and the Use Committee (IACUC APN# 2015-100932). Fresh tumour pieces derived from a RCC patient were implanted orthotopically under the kidney capsule of 4-to-6 weeks-old non-obese diabetic severe combined immunodeficient (NOD/SCID) mice as previously described^53,54^. Tumour growth was monitored weekly by physical palpation. Once tumours reached approximately ∼7mm, the mice were euthanized.

Establishment of melanoma patient derived xenografts were performed as previously described^55^. 100 cells in a final volume of 50μL were subcutaneously injected into the right flank of NOD.CB17-*Prkdc^scid^ Il2rg^tm1Wjl^*/SzJ (NSG) mice, and subcutaneous tumours were measured weekly with calipers until tumours reached 2.5 cm in its largest diameter.

### Human tissue samples

Patients were recruited to a UTSW IRB-approved study (STU2019-1061, NCT04623502) and informed consent was obtained. ccRCC and adjacent kidney tissues were snap frozen in liquid nitrogen immediately after sampling with the Attending Pathologist or Pathology Assistant. Tissue histology was confirmed with hematoxylin and eosin (H&E) staining of sections flanking the snap frozen sampled tissue.

### Statistics and reproducibility

GraphPad PRISM v.7 and Microsoft Excel v.15.21.1 software were used for statistical analysis. Analyst 1.6.3 software (SCIEX) was used for metabolomic analyses. Error bars, *P* values and statistical tests are reported in the figure captions. All in vitro experiments were performed at least thrice with similar results. All mouse experiments were performed at least twice with similar results. Patient-derived xenografts and human sample analysis were done once due to limiting amounts of patient tissue. Both technical and biological replicates were reliably reproduced.

## ACKNOWLEDGEMENTS

We thank Sean Morrison for kindly providing melanoma PDXs. The KP mouse cancer cell lines HY15549 and Tom2 are gifts from N. Bardeesy and T. Papagiannakopoulos. J.G.B. was supported by NCI (4R00CA248838-02) and the Cancer Prevention and Research Institute of Texas (CPRIT RR210059). R.J.D. is supported by the Howard Hughes Medical Institute Investigator Program and grants from the NIH (R35CA220449, P50CA196516). D.B. was supported by grants from the NIH (F31CA239330). K.B. is supported by the NIH/NIDDK (R01 DK123323-01) and a Mark Foundation Emerging Leader Award and is a Searle and Pew-Stewart Scholar.

ccRCC PDX experiments were supported by a National Institutes of Health Kidney Specialized Programs of Research Excellence (SPORE) (P50CA196516). V.T.T. and J.B. are supported by a National Institutes of Health Kidney Specialized Programs of Research Excellence (SPORE) (P50CA196516). We acknowledge the assistance of the University of Texas Southwestern Tissue Management Shared Resource, a shared resource at the Simmons Comprehensive Cancer Center, which is supported in part by the National Cancer Institute under award number P30CA142543. We thank the patients that provided their tissue for research.

R.J.W. is supported by NIGMS R35-GM150736 and NHLBI R21-HL167091. We would like to thank Parastoo Azadi and the UGA Analytical Services & Training Core for providing instrumentation to perform the glycosaminoglycan analyses.

## CONTRIBUTIONS

J.G.B. and D.C. conceived the project and designed the experiments with input from K.B. D.C., L.S. and N.K. performed most of the experiments with assistance of D.B., S-C.H., A.P., and A.P.S. R.J.W and A.B. performed mass spectrometry analyses of glycosaminoglycans. K.L. and L.C. performed computational analyses. V.T.T., D.L.C. and J.B provided patient-derived xenograft fresh tissues. V.M., R.J.D., D.B. and B.B. provided human ccRCC and adjacent kidney specimens. F.C. performed metabolomic analysis. J.G.B. wrote the manuscript with help from K.B., R.J.D. and D.C. All authors reviewed the manuscript.

## EXTENDED DATA FIGURE LEGENDS

**Extended Data Figure 1.**
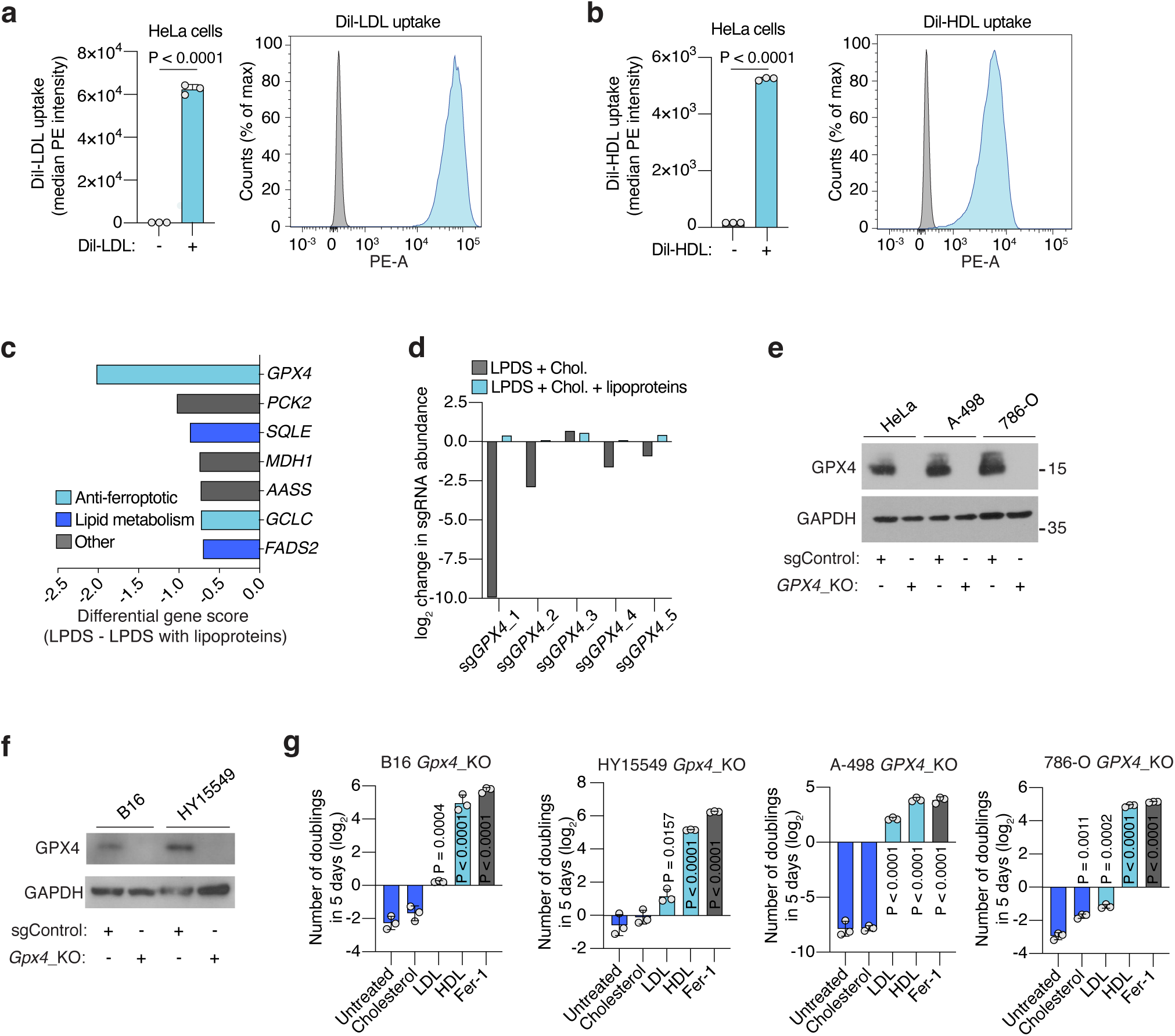
GPX4 loss is compensated by lipoprotein supplementation. **a.** Cellular uptake of DiI-LDL in HeLa cells measured as median PE intensity after incubation or not with DiI-LDL (left). Representative flow cytometry plot of HeLa cells treated (blue) or not (grey) with DiI-LDL (right). **b.** Cellular uptake of DiI-HDL in HeLa cells measured as median PE intensity after incubation or not with DiI-HDL (left). Representative flow cytometry plot of HeLa cells treated (blue) or not (grey) with DiI-HDL (right). **c.** Plot of differential gene scores in HeLa cells supplemented with lipoproteins (LPDS + lipoproteins) or not (LPDS). Anti-ferroptotic genes are shown in light blue and lipid metabolism genes are shown in dark blue. LPDS: lipoprotein-depleted serum. **d.** Individual sgRNA scores (log_2_) for *GPX4* under indicated conditions. **e.** Immunoblot analysis of human GPX4 sgControl cells or *GPX4*_KO isogenic lines. GAPDH is included as a loading control. **f.** Immunoblot analysis of mouse GPX4 sgControl cells or *Gpx4*_KO isogenic lines. GAPDH is included as a loading control. **g.** Number of doublings (log_2_) in 5 days of indicated GPX4-deficient cell lines in vitro either untreated or supplemented with cholesterol (5 μg/mL), LDL (50 μg/mL), HDL (50 μg/mL) or Fer-1 (1 μM). **A, b, g,** Bars represent mean ± s.d.; **a, b, g,** n=3 biological replicates. Statistical significance determined by two-tailed unpaired t-tests as indicated or compared to untreated cells (**g**).

**Extended Data Figure 2.**
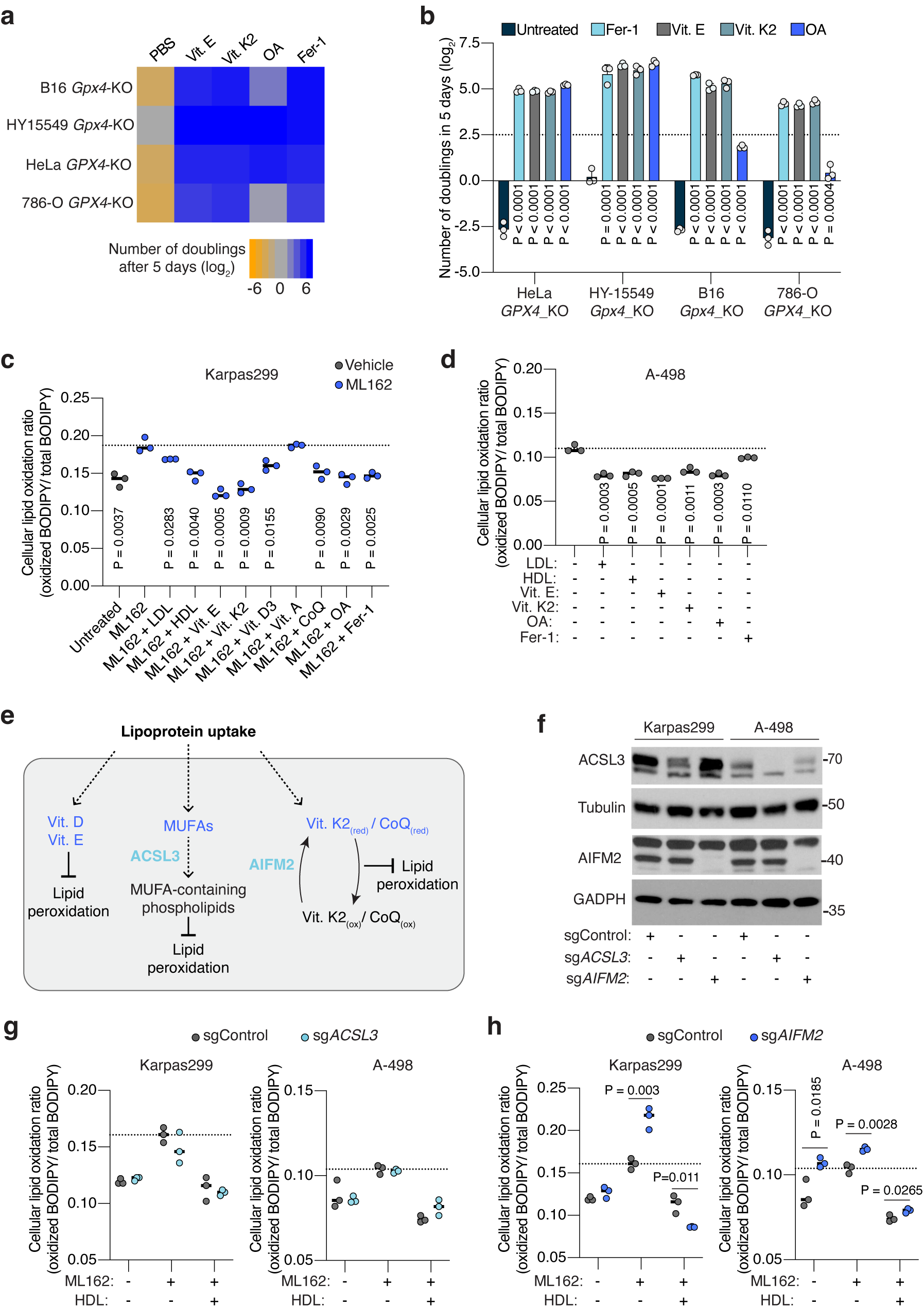
Lipoproteins carry multiple lipid species that could inhibit lipid peroxidation. **a.** Heatmap showing the number of doublings (log_2_) in 5 days of mouse (*Gpx4*) or human (*GPX4*) knockout cell lines in cell culture under the supplementation of PBS, vitamin E (Vit. E, 10 μM), vitamin K2 (Vit. K2, 10 μM), oleic acid (OA, 250 μM) or ferrostatin-1 (Fer-1, 1 μM). **b.** Number of doublings (log_2_) in 5 days of B16 and HY15549 *Gpx4*-*KO* and HeLa and 786-O *GPX4*-KO cells in vitro either untreated or supplemented with Fer-1 (1 μM), vitamin E (10 μM), vitamin K2 (10 μM), or OA (250 μM). **c.** Cellular lipid oxidation ratio using BODIPY-C11 in HeLa cells in the absence (gray) or presence (blue) of a GPX4 inhibitor (ML162, 125 nM) under supplementation or not of LDL (50 μg/mL), HDL (50 μg/mL), vitamin E (10 μM), vitamin K2 (10 μM), vitamin D3 (10 μM), vitamin A (10 μM), CoQ10 (10 μM), OA (250 μM) or Fer-1 (1 μM). **d.** Cellular lipid oxidation ratio using BODIPY-C11 in A-498 cells with intact GPX4 activity and under supplementation or not of LDL (50 μg/mL), HDL (50 μg/mL), vitamin E (10 μM), vitamin K2 (10 μM), OA (250 μM) or Fer-1 (1 μM). **e.** Scheme highlighting genes (light blue) that affect the cellular ability to utilize lipoprotein-transported lipids (dark blue) that inhibit lipid peroxidation. **f.** Immunoblot analysis of ACSL3 and AIFM2 in the indicated cell lines transduced with a sgControl, sg*ACSL3* or sg*AIFM2*. GAPDH is included as a loading control. **g.** Cellular lipid oxidation ratio using BODIPY-C11 in the indicated Karpas299 (left) or A-498 (right) cell lines transduced with sg*ACSL3* (light blue) or with a sgControl (grey) under GPX4 inhibition (ML162: 75 nM for Karpas299, 175 nM for A-498) and HDL supplementation (50 μg/mL). **h.** Cellular lipid oxidation ratio using BODIPY-C11 in the indicated Karpas299 (left) or A-498 (right) cell lines transduced with sg*AIFM2* (dark blue) or with a sgControl (grey) under GPX4 inhibition (ML162: 75 nM for Karpas299, 175 nM for A-498) and HDL supplementation (50 μg/mL). **b,** Bars represent mean ± s.d.; **c, d, g, h,** bars represent median; **b, c, d, g, h,** n=3 biologically independent samples. Statistical significance determined by two-tailed unpaired t-tests as indicated or compared to untreated cells (**b, d**) or ML162 treated cells (**c**).

**Extended Data Figure 3.**
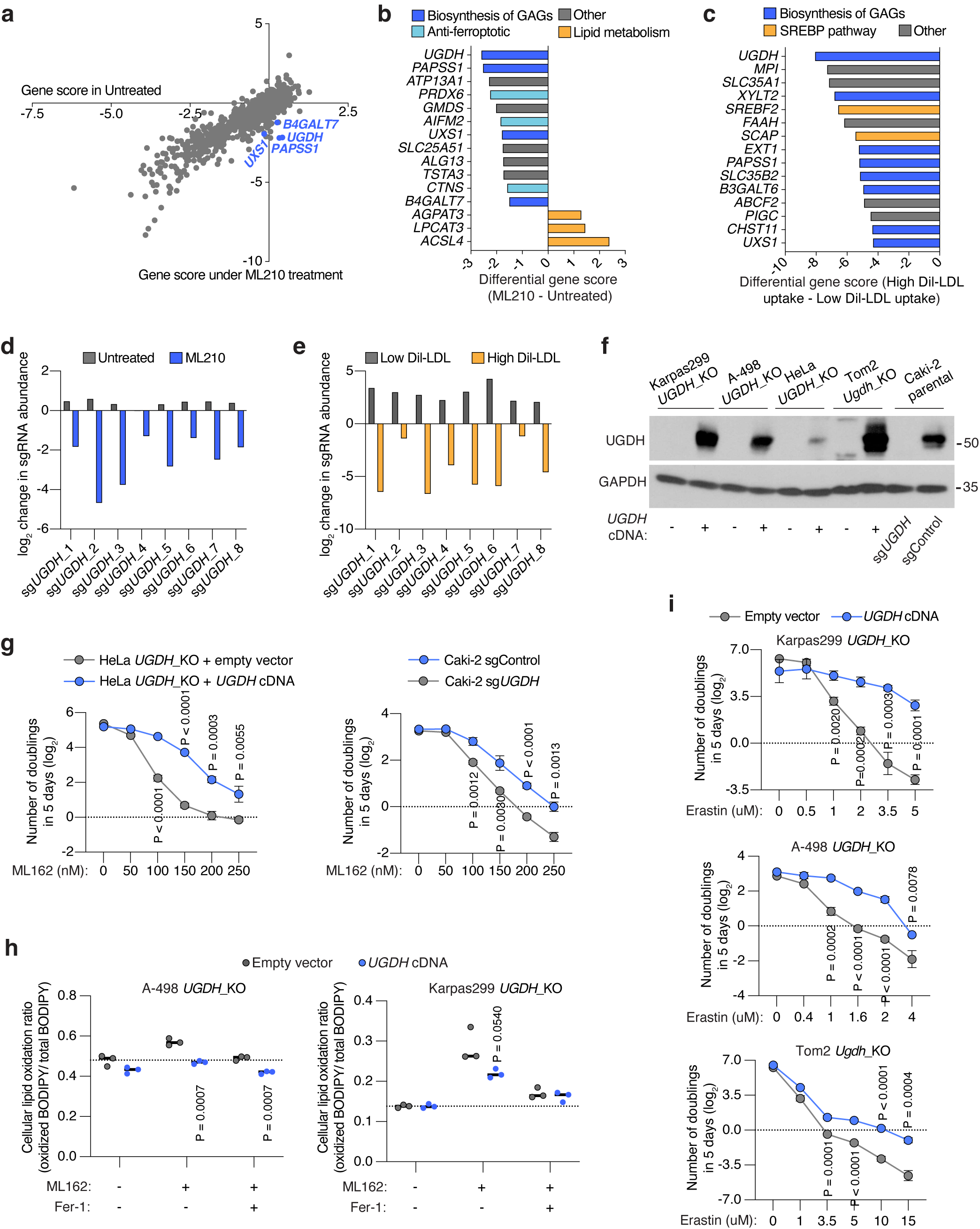
Loss of UGDH sensitizes cancer cells to ferroptosis in vitro. **a.** Gene essentiality plot showing genes scores in untreated (x-axis) and ML210-treated (GPX4 inhibition) Karpas299 cells. Glycosaminoglycan (GAG) biosynthesis genes are highlighted (dark blue). **b.** Top-scoring genes under ML210 treatment. Negative scores represent genes whose loss potentiates ML210 toxicity; positive scores represent genes whose loss provides resistance to ML210. GAG biosynthesis genes are shown in dark blue, canonical anti-ferroptotic genes are shown in light blue, and lipid metabolism genes are shown in yellow. **c.** Top-scoring genes essential for DiI-LDL uptake. Negative scores represent genes whose loss reduce cellular DiI-LDL uptake. GAG biosynthesis genes are shown in dark blue and genes in canonical lipoprotein uptake pathways highlighted in yellow. **d.** Individual sgRNA scores for *UGDH* in untreated and ML210-treated conditions. **e.** Individual sgRNA scores for *UGDH* in high and low DiI-LDL uptake populations. **f.** Immunoblot analysis of UGDH in the indicated isogenic cell lines transduced with sg*UGDH* and expressing or not a sgRNA-resistant *UGDH* cDNA. GAPDH is included as a loading control. **g.** Number of doublings (log_2_) in 5 days of indicated HeLa (left) or Caki-2 (right) cell lines expressing UGDH (blue) or not (grey) under the indicated concentrations of the GPX4 inhibitor ML162. **h.** Cellular lipid oxidation ratio using BODIPY-C11 in the indicated A-498 (left) or Karpas299 (right) *UGDH*_KO cell lines expressing a sgRNA resistant *UGDH* cDNA (blue) or an empty vector (grey) under the indicated conditions of GPX4 inhibition (ML162: 200 nM for Karpas299, 250 nM for A-498) and Fer-1 supplementation (1μM). **i.** Number of doublings (log_2_) in 5 days of the indicated Karpas299 (top), A-498 (middle) and Tom2 (bottom) UGDH-deficient cell lines expressing a sgRNA resistant *UGDH* cDNA (dark blue) or an empty vector (grey) under the indicated concentrations of erastin. **g-i,** Bars represent mean ± s.d.; **g-i,** n=3 biologically independent samples. Statistical significance determined by two-tailed unpaired t-tests compared to empty vector transduced cells.

**Extended Data Figure 4.**
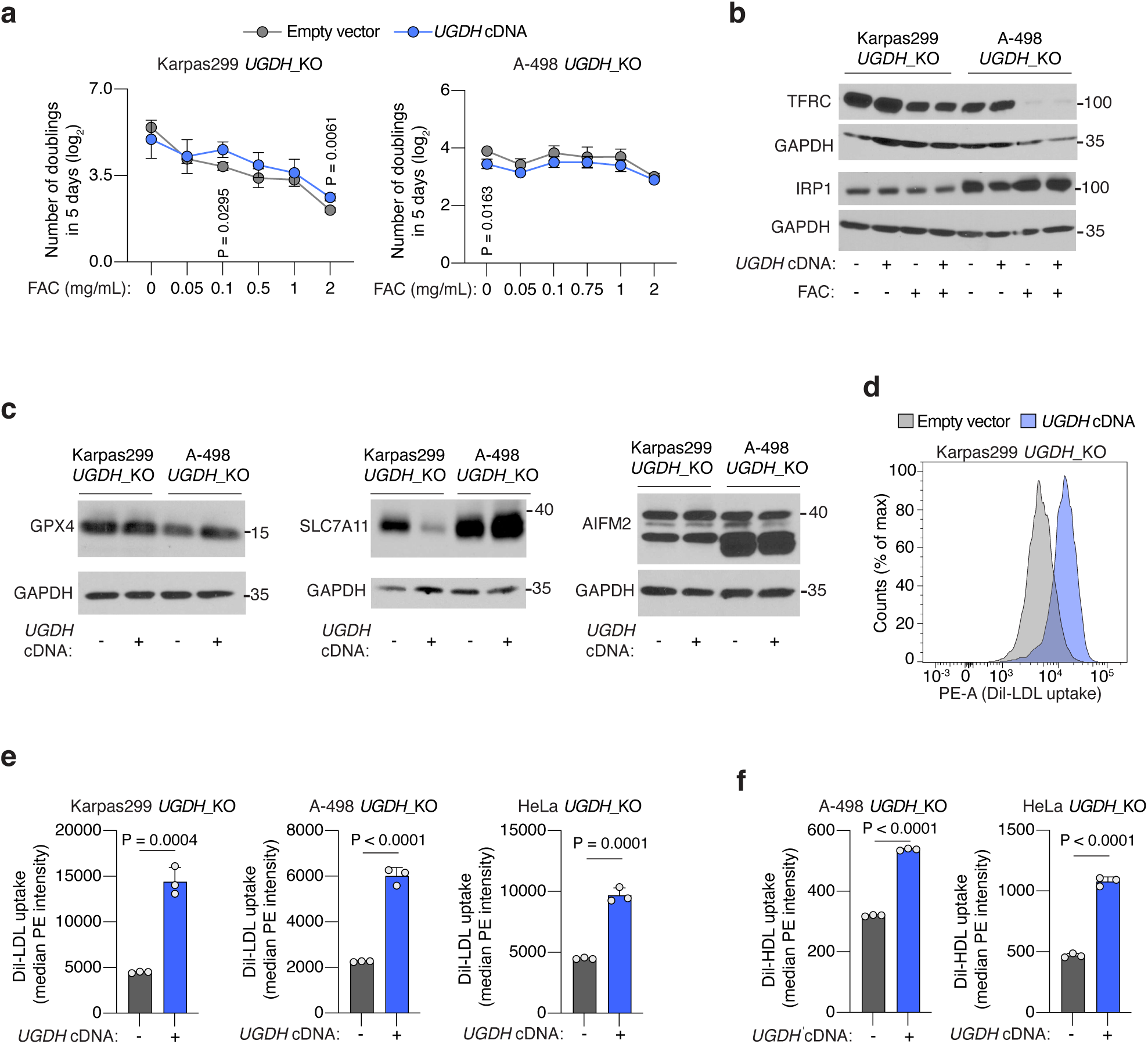
UGDH is a major determinant of LDL and HDL uptake in cancer cells. **a.** Number of doublings (log_2_) in 5 days of the indicated Karpas299 (left) and A-498 (right) *UGDH*_KO cell lines expressing a sgRNA resistant *UGDH* cDNA (blue) or an empty vector (grey) under the indicated concentrations of ferrous ammonium citrate (FAC). **b.** Immunoblot analysis of TFRC (top) and IRP1 (bottom) in the indicated *UGDH*_KO cell lines expressing a sgRNA resistant *UGDH* cDNA or an empty vector treated or not with FAC (0.1 mg/mL). GAPDH is included as a loading control. **c.** Immunoblot analysis of GPX4 (left), SLC7A11 (center), and AIFM2 (right) in the indicated *UGDH*_KO cell lines expressing a sgRNA resistant *UGDH* cDNA or an empty vector. GAPDH is included as a loading control. **d.** Representative flow cytometry plot showing the uptake of DiI-LDL (PE median intensity) in Karpas299 *UGDH*_KO cells expressing a sgRNA resistant *UGDH* cDNA (blue) or an empty vector (grey). **e.** Cellular uptake of DiI-LDL measured as median PE intensity in the indicated Karpas299 (left), A-498 (center) and HeLa (right) *UGDH*_KO cells expressing a sgRNA resistant *UGDH* cDNA (blue) or an empty vector (grey). **f.** Cellular uptake of DiI-HDL measured as median PE intensity in the indicated A-498 (left) and HeLa (right) *UGDH*_KO cells expressing a sgRNA resistant *UGDH* cDNA (blue) or an empty vector (grey). **a, e, f,** Lines or bars represent mean ± s.d.; **a, e, f,** n=3 biologically independent samples. Statistical significance determined by two-tailed unpaired t-tests.

**Extended Data Figure 5.**
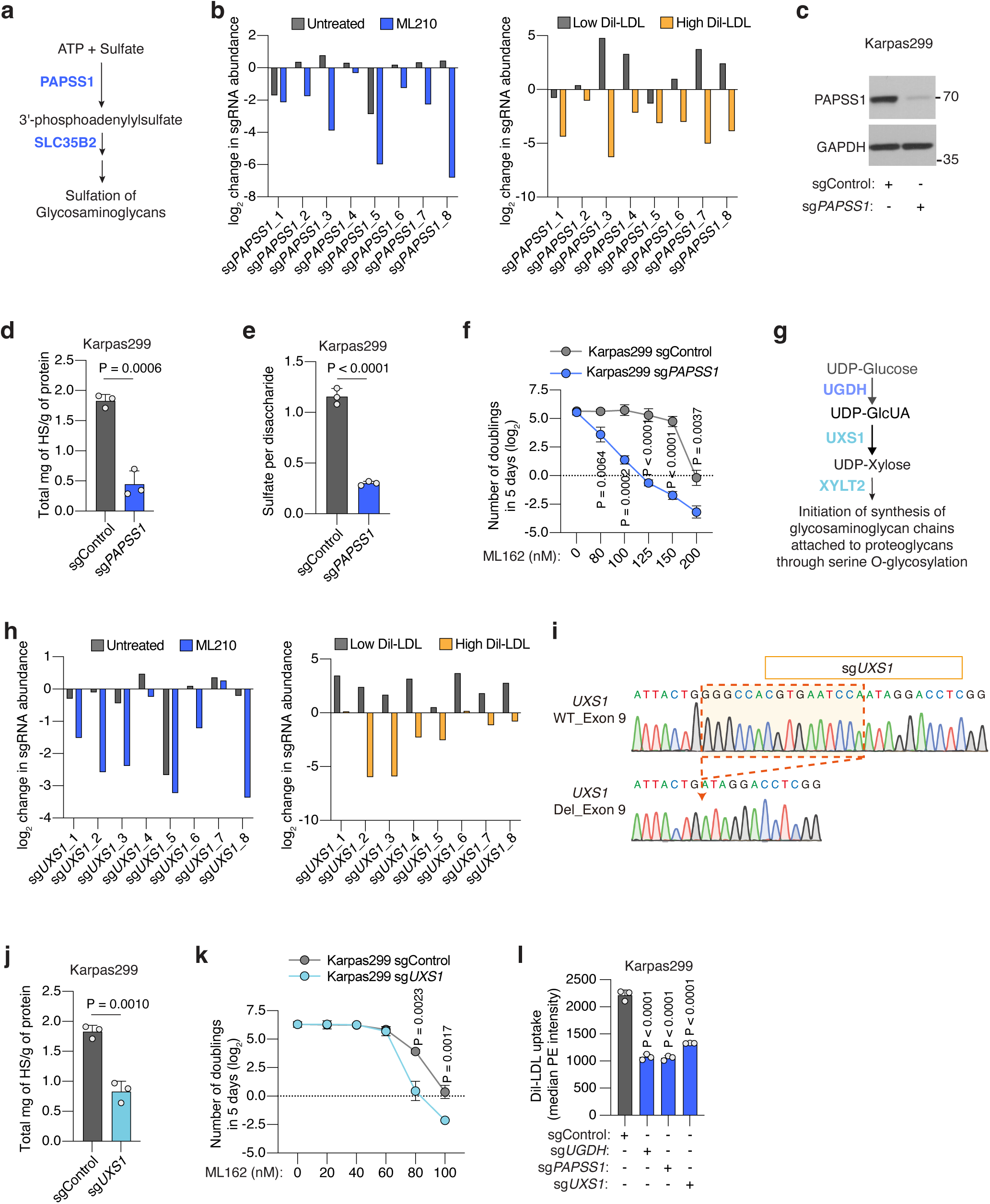
Sulfation of GAGs and xylose synthesis is essential for lymphoma cells to resist ferroptosis and take up lipoproteins. **a.** Sulfation of glycosaminoglycans relies on the production of 3’-phosphoadenylylsulfate by *PAPSS1*, and on its transports into the Golgi by *SLC35B2*. Both genes (blue) scored in our screens. **b.** Individual sgRNA scores for *PAPSS1* in the presence and absence of ML210 (left), or in high and low DiI-LDL populations (right). **c.** Immunoblot analysis of PAPSS1 in Karpas299 cells transduced with a sgControl (grey) or sg*PAPSS1* (blue). GAPDH is included as a loading control. **d.** Quantification of total milligrams of heparan sulfate (HS) per grams of protein in Karpas299 cells transduced with a sgControl (grey) or sg*PAPSS1* (blue). **e.** Quantification of total sulfate per glycosaminoglycan disaccharide in Karpas299 cells transduced with a sgControl (grey) or sg*PAPSS1* (blue). **f.** Number of doublings (log_2_) in 5 days of Karpas299 cells transduced with a sgControl (grey) or sg*PAPSS1* (blue) under the indicated concentrations of the GPX4 inhibitor ML162. **g.** Glucuronic acid (GlcUA), formed by UGDH, is converted to UDP-Xylose via *UXS1*. UDP-Xylose is the first monosaccharide used in the initiation of O-glycosylation that attaches GAG chains to serine-residues of proteoglycans. Genes involved in this process (blue) scored in our screens. **h.** Individual sgRNA scores for *UXS1* in the presence and absence of ML210 (left), or in high and low DiI-LDL populations (right). **i.** Sanger sequencing of *UXS1* gene exon 9 in Karpas299 parental cells (WT) or transduced with sg*UXS1*. **j.** Quantification of total milligrams of heparan sulfate (HS) per grams of protein in Karpas299 cells transduced with a sgControl (grey) or sg*UXS1* (blue). **k.** Number of doublings (log_2_) in 5 days of Karpas299 cells transduced with a sgControl (grey) or sg*UXS1* (blue) under the indicated concentrations of the GPX4 inhibitor ML162. **i.** Cellular uptake of DiI-LDL measured as median PE intensity in the indicated Karpas299 cells transduced with a sgControl, sg*UGDH*, sg*PAPSS1* or sg*UXS1* assessed by flow cytometry. **d-f, j-l,** Bars or lines represent mean ± s.d.; **d-f, j-l,** n=3 biologically independent samples. Statistical significance determined by two-tailed unpaired t-tests as indicated or compared to sgControl cells (**l**).

**Extended Data Figure 6.**
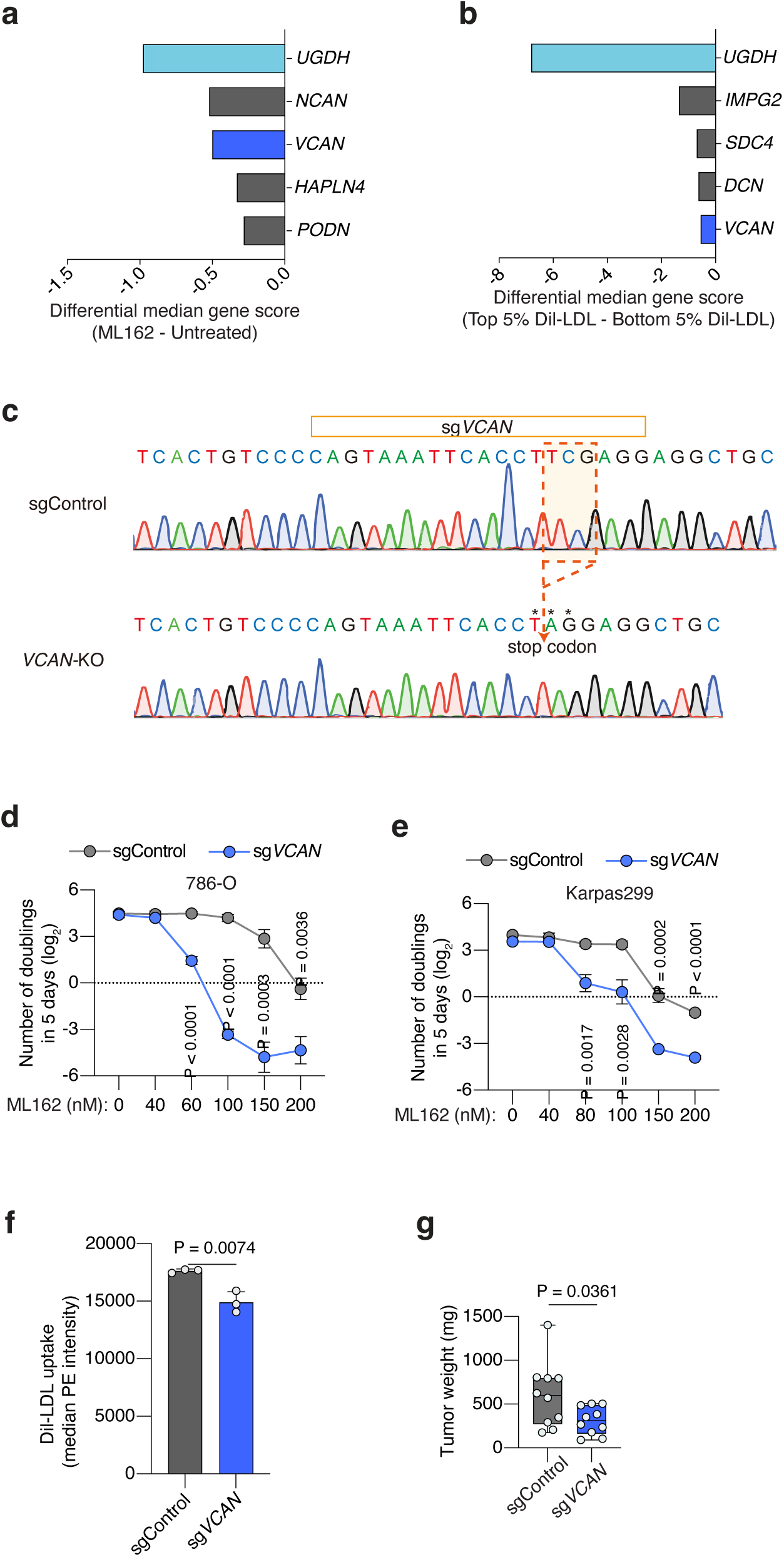
The proteoglycan VCAN modestly increases resistance to ferroptosis and lipoprotein uptake in cancer cells. **a.** Plot of differential gene scores under ML162 treatment relative to untreated cells. Negative scores represent genes whose loss potentiates ML162 toxicity. *UGDH* was included as a positive control (light blue). *VCAN* (blue) was a common hit between the two screens. **b.** Plot of differential gene scores in high DiI-LDL uptake population compared to low DiI-LDL cells. Negative scores represent genes whose loss reduce cellular DiI-LDL uptake. *UGDH* was included as a positive control (light blue) and *VCAN* (blue) was a common hit between the two screens. **c.** Sanger sequencing of *VCAN* gene exon 9 in Karpas299 parental cells (WT) or transduced with sg*UXS1*. **d.** Number of doublings (log_2_) in 5 days of 786-O transduced with a sgControl (grey) or sg*VCAN* (blue) under the indicated concentrations of the GPX4 inhibitor ML162. **e.** Number of doublings (log_2_) in 5 days of Karpas299 cells transduced with a sgControl (grey) or sg*VCAN* (dark blue) under the indicated concentrations of the GPX4 inhibitor ML162. **f.** Cellular uptake of DiI-LDL measured as median PE intensity in the indicated Karpas299 cells transduced with a sgControl (grey) or sg*VCAN* (blue) assessed by flow cytometry. **g.** Tumour weight resulting from implantation of Karpas299 cells transduced with a sgControl (grey) or sg*VCAN* (blue) in 6-12 weeks old immunodeficient mice. **d-g,** Bars or lines represent mean ± s.d.; **d-g,** n=3 biologically independent samples. Statistical significance determined by two-tailed unpaired t-tests.

**Extended Data Figure 7.**
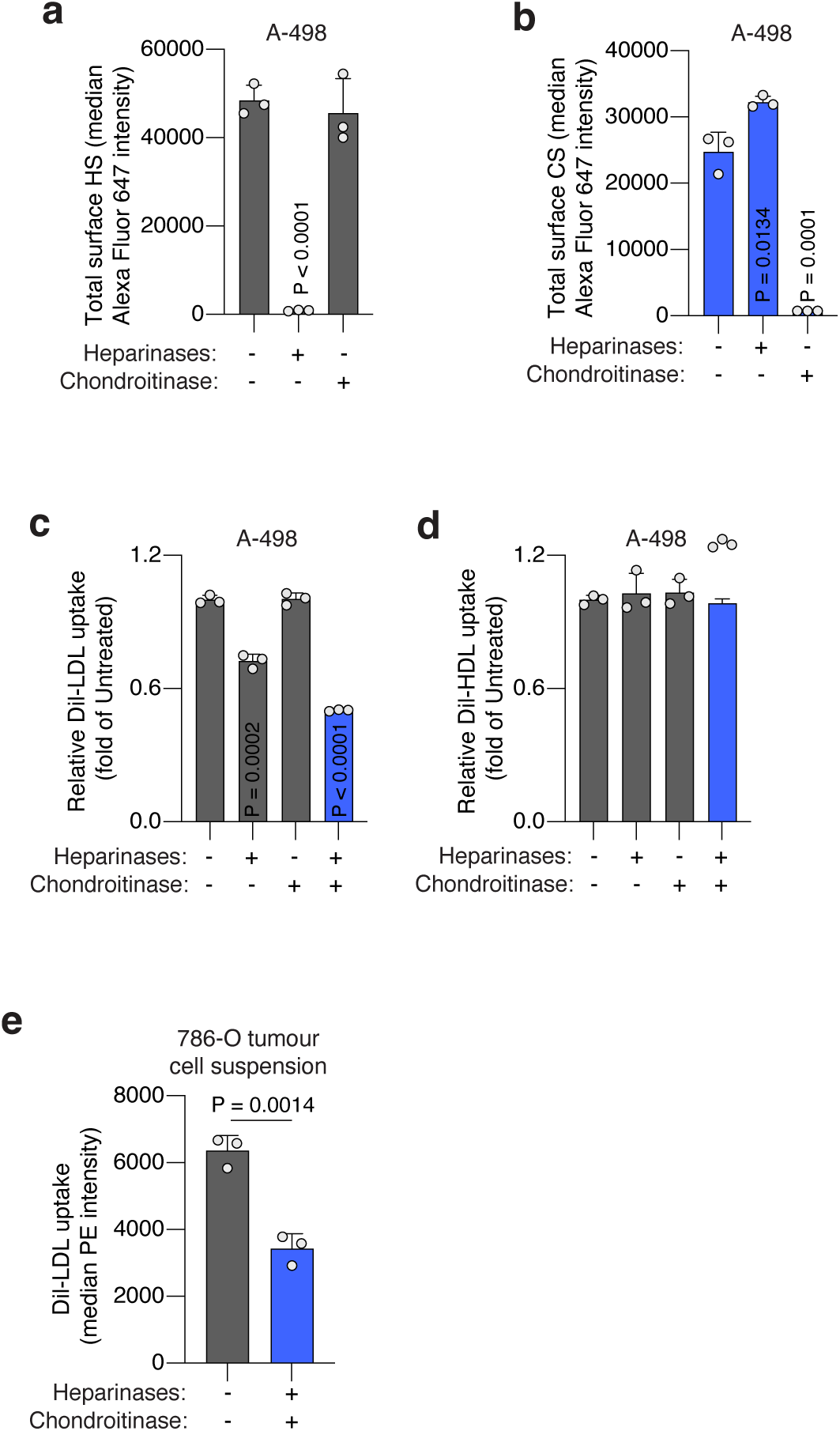
Cell-surface sulfated glycosaminoglycans promote lipoprotein uptake. **a.** Total cell surface HS measured as median Alexa Fluor 647 intensity in A-498 cells treated or not with heparinases (0.1 U/mL) or chondroitinases (0.1 U/mL) assessed by flow cytometry. **b.** Total cell surface CS measured as median Alexa Fluor 647 intensity in A-498 cells treated or not with heparinases (0.1 U/mL) or chondroitinases (0.1 U/mL) assessed by flow cytometry. **c.** Fold change in DiI-LDL uptake of A-498 cells upon treatment with heparinases (0.1 U/mL), chondroitinases (0.1 U/mL), or both (blue), relative to uptake of untreated cells assessed by flow cytometry. **d.** Fold change in DiI-HDL uptake of A-498 cells upon treatment with heparinases (0.1 U/mL), chondroitinases (0.1 U/mL), or both (blue), relative to uptake of untreated cells, assessed by flow cytometry. **e.** Uptake of DiI-LDL measured as median PE intensity in a cell suspension of 786-O xenograft tumour fresh tissue resected from mice left untreated (grey) or after combined treatment with heparinases (1.0 U/mL) and chondroitinases (1.0 U/mL) (blue) assessed by flow cytometry. **a-e,** Bars represent mean ± s.d.; **a-e,** n=3 biologically independent samples. Statistical significance determined by two-tailed unpaired t-tests as indicated or compared to untreated cells (**a-d**).

**Extended Data Figure 8.**
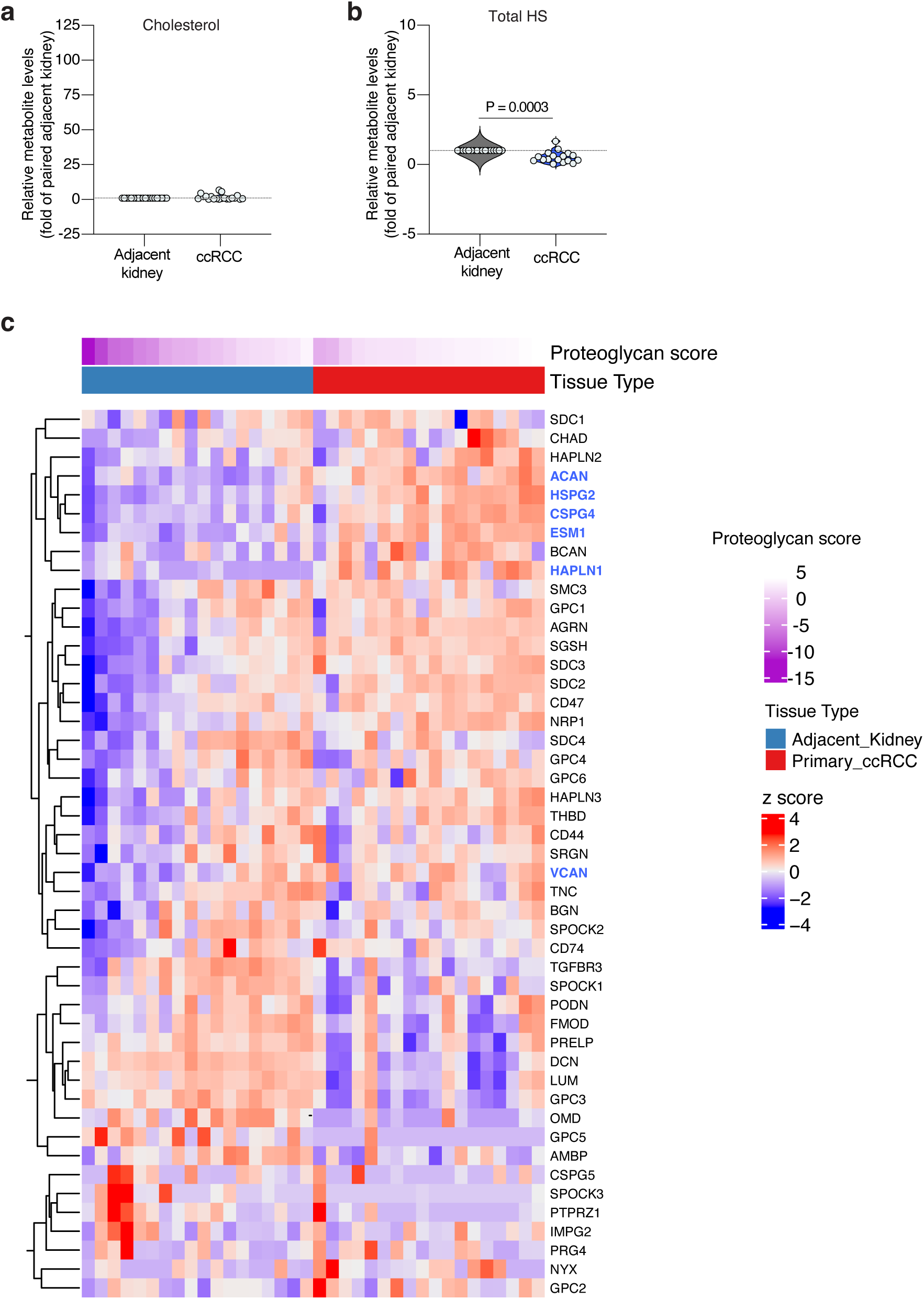
Proteoglycan expression profile in human clear cell renal cell carcinomas (ccRCCs) and adjacent kidney. **a.** Violin plot showing the relative levels of cholesterol in ccRCC patient tissues (blue) compared to paired adjacent kidney (grey). **b.** Violin plot showing the relative levels of total heparan sulfate (HS) per gram of protein in ccRCC patient tissues (blue) compared to paired adjacent kidney (grey). **c.** Heatmap showing expression of individual proteoglycan genes in ccRCC tumours and paired adjacent kidney. Each sample was assigned a proteoglycan score, based on their expression of all proteoglycans (top). Relevant proteoglycans scoring are highlighted in blue. **a, b,** n=17-20 biologically independent samples. Statistical significance was determined by a two-tailed unpaired t-test.

**Extended Data Figure 9.**
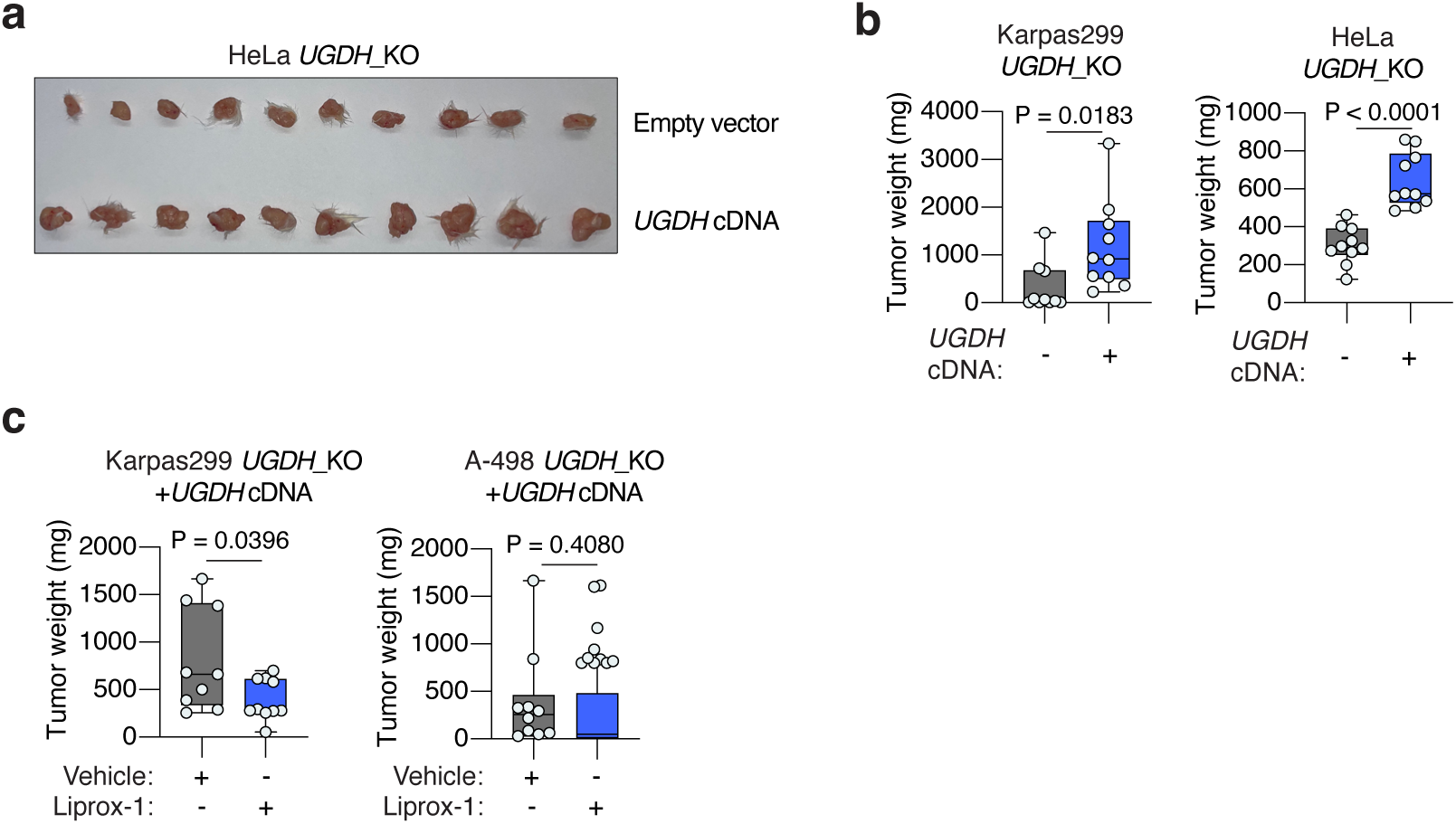
The anti-ferroptotic effect of UGDH is essential for tumour growth. **a.** Representative tumour images of HeLa *UGDH*_KO cells expressing a sgRNA resistant *UGDH* cDNA or an empty vector, and implanted subcutaneously in 6-12 weeks old immunodeficient mice. **b.** Tumour weight resulting from implantation of the indicated *UGDH*_KO cell lines in immunodeficient mice. **c.** Tumour weights resulting from implantation of the indicated *UGDH*_KO cell lines expressing a sgRNA-resistant *UGDH* cDNA on immunodeficient mice treated with vehicle (grey) or Lip-1 (blue) through daily intraperitoneal injection. **b, c,** Boxes represent the median, and the first and third quartiles, and the whiskers represent the minimum and maximum of all data points.; **b, c,** n=10 biologically independent samples. Statistical significance determined by two-tailed unpaired t-tests.

## REFERENCES

1. Medes, G., Thomas, A. & Weinhouse, S. Metabolism of neoplastic tissue. IV. A study of lipid synthesis in neoplastic tissue slices in vitro. Cancer Res 13, 27–29 (1953).

2. Mashima, T., Seimiya, H. & Tsuruo, T. De novo fatty-acid synthesis and related pathways as molecular targets for cancer therapy. Br. J. Cancer 100, 1369–1372 (2009).

3. Garcia-Bermudez, J. et al. Squalene accumulation in cholesterol auxotrophic lymphomas prevents oxidative cell death. Nature 567, 118–122 (2019).

4. Riscal, R. et al. Cholesterol Auxotrophy as a Targetable Vulnerability in Clear Cell Renal Cell Carcinoma. Cancer Discov 11, 3106–3125 (2021).

5. Zhang, M. et al. Adipocyte-Derived Lipids Mediate Melanoma Progression via FATP Proteins. Cancer Discov 8, 1006–1025 (2018).

6. Villa, G. R. et al. An LXR-Cholesterol Axis Creates a Metabolic Co-Dependency for Brain Cancers. Cancer Cell 30, 683–693 (2016).

7. Dixon, S. J. et al. Ferroptosis: an iron-dependent form of nonapoptotic cell death. Cell 149, 1060–1072 (2012).

8. Lang, X. et al. Radiotherapy and Immunotherapy Promote Tumoral Lipid Oxidation and Ferroptosis via Synergistic Repression of SLC7A11. Cancer Discov 9, 1673–1685 (2019).

9. Ubellacker, J. M. et al. Lymph protects metastasizing melanoma cells from ferroptosis. Nature 585, 113–118 (2020).

10. Yang, W. S. et al. Regulation of ferroptotic cancer cell death by GPX4. Cell 156, 317–331 (2014).

11. Freitas, F. P. et al. 7-Dehydrocholesterol is an endogenous suppressor of ferroptosis. Nature 626, 401–410 (2024).

12. Traber, M. G. & Atkinson, J. Vitamin E, antioxidant and nothing more. Free Radic Biol Med 43, 4–15 (2007).

13. Mishima, E. et al. A non-canonical vitamin K cycle is a potent ferroptosis suppressor. Nature 608, 778–783 (2022).

14. Bersuker, K. et al. The CoQ oxidoreductase FSP1 acts parallel to GPX4 to inhibit ferroptosis. Nature 575, 688–692 (2019).

15. Doll, S. et al. FSP1 is a glutathione-independent ferroptosis suppressor. Nature 575, 693–698 (2019).

16. Magtanong, L. et al. Exogenous Monounsaturated Fatty Acids Promote a Ferroptosis-Resistant Cell State. Cell Chem Biol 26, 420–432.e9 (2019).

17. Quehenberger, O. & Dennis, E. A. The human plasma lipidome. N Engl J Med 365, 1812–1823 (2011).

18. Rigotti, A. Absorption, transport, and tissue delivery of vitamin E. Mol Aspects Med 28, 423–436 (2007).

19. Birsoy, K. et al. An Essential Role of the Mitochondrial Electron Transport Chain in Cell Proliferation Is to Enable Aspartate Synthesis. Cell 162, 540–551 (2015).

20. Lu, B. et al. Identification of PRDX6 as a regulator of ferroptosis. Acta Pharmacol Sin 40, 1334– 1342 (2019).

21. Matsuda, M. et al. SREBP cleavage-activating protein (SCAP) is required for increased lipid synthesis in liver induced by cholesterol deprivation and insulin elevation. Genes Dev 15, 1206–1216 (2001).

22. Pomin, V. H. & Mulloy, B. Glycosaminoglycans and Proteoglycans. Pharmaceuticals (Basel) 11, 27 (2018).

23. Sasisekharan, R., Shriver, Z., Venkataraman, G. & Narayanasami, U. Roles of heparan-sulphate glycosaminoglycans in cancer. Nat Rev Cancer 2, 521–528 (2002).

24. Asimakopoulou, A. P., Theocharis, A. D., Tzanakakis, G. N. & Karamanos, N. K. The biological role of chondroitin sulfate in cancer and chondroitin-based anticancer agents. In Vivo 22, 385–389 (2008).

25. Yang, W. S. & Stockwell, B. R. Synthetic lethal screening identifies compounds activating iron-dependent, nonapoptotic cell death in oncogenic-RAS-harboring cancer cells. Chem Biol 15, 234–245 (2008).

26. Volpi, N. & Tarugi, P. Influence of chondroitin sulfate charge density, sulfate group position, and molecular mass on Cu2+-mediated oxidation of human low-density lipoproteins: effect of normal human plasma-derived chondroitin sulfate. J Biochem 125, 297–304 (1999).

27. Koppula, P., Zhang, Y., Shi, J., Li, W. & Gan, B. The glutamate/cystine antiporter SLC7A11/xCT enhances cancer cell dependency on glucose by exporting glutamate. J Biol Chem 292, 14240–14249 (2017).

28. Flood, C. et al. Identification of the proteoglycan binding site in apolipoprotein B48. J Biol Chem 277, 32228–32233 (2002).

29. Goldstein, J. L., Basu, S. K., Brunschede, G. Y. & Brown, M. S. Release of low density lipoprotein from its cell surface receptor by sulfated glycosaminoglycans. Cell 7, 85–95 (1976).

30. Brown, M. S., Kovanen, P. T. & Goldstein, J. L. Regulation of plasma cholesterol by lipoprotein receptors. Science 212, 628–635 (1981).

31. Pownall, H. J., Rosales, C., Gillard, B. K. & Gotto, A. M. High-density lipoproteins, reverse cholesterol transport and atherogenesis. Nat Rev Cardiol 18, 712–723 (2021).

32. Klaassen, C. D. & Boles, J. W. Sulfation and sulfotransferases 5: the importance of 3’-phosphoadenosine 5’-phosphosulfate (PAPS) in the regulation of sulfation. FASEB J 11, 404–418 (1997).

33. Kearns, A. E., Campbell, S. C., Westley, J. & Schwartz, N. B. Initiation of chondroitin sulfate biosynthesis: a kinetic analysis of UDP-D-xylose: core protein beta-D-xylosyltransferase. Biochemistry 30, 7477–7483 (1991).

34. Basu, A., Patel, N. G., Nicholson, E. D. & Weiss, R. J. Spatiotemporal diversity and regulation of glycosaminoglycans in cell homeostasis and human disease. Am J Physiol Cell Physiol 322, C849– C864 (2022).

35. Wight, T. N. Versican: a versatile extracellular matrix proteoglycan in cell biology. Curr Opin Cell Biol 14, 617–623 (2002).

36. Nandadasa, S. et al. The versican-hyaluronan complex provides an essential extracellular matrix niche for Flk1+ hematoendothelial progenitors. Matrix Biol 97, 40–57 (2021).

37. Noborn, F. et al. Site-specific identification of heparan and chondroitin sulfate glycosaminoglycans in hybrid proteoglycans. Sci Rep 6, 34537 (2016).

38. Qi, X., Li, Q., Che, X., Wang, Q. & Wu, G. The Uniqueness of Clear Cell Renal Cell Carcinoma: Summary of the Process and Abnormality of Glucose Metabolism and Lipid Metabolism in ccRCC. Front Oncol 11, 727778 (2021).

39. Gatto, F. et al. Glycosaminoglycan Profiling in Patients’ Plasma and Urine Predicts the Occurrence of Metastatic Clear Cell Renal Cell Carcinoma. Cell Rep 15, 1822–1836 (2016).

40. Evanko, S. P. et al. A Role for HAPLN1 During Phenotypic Modulation of Human Lung Fibroblasts In Vitro. J Histochem Cytochem 68, 797–811 (2020).

41. Xu, Y.-X. et al. The glycosylation-dependent interaction of perlecan core protein with LDL: implications for atherosclerosis. J Lipid Res 56, 266–276 (2015).

42. Piskounova, E. et al. Oxidative stress inhibits distant metastasis by human melanoma cells. Nature 527, 186–191 (2015).

43. Zilka, O. et al. On the Mechanism of Cytoprotection by Ferrostatin-1 and Liproxstatin-1 and the Role of Lipid Peroxidation in Ferroptotic Cell Death. ACS Cent Sci 3, 232–243 (2017).

44. Gonzales, J. C., Gordts, P. L. S. M., Foley, E. M. & Esko, J. D. Apolipoproteins E and AV mediate lipoprotein clearance by hepatic proteoglycans. J Clin Invest 123, 2742–2751 (2013).

45. MacArthur, J. M. et al. Liver heparan sulfate proteoglycans mediate clearance of triglyceride-rich lipoproteins independently of LDL receptor family members. J Clin Invest 117, 153–164 (2007).

46. Gordts, P. L. S. M. et al. Reducing macrophage proteoglycan sulfation increases atherosclerosis and obesity through enhanced type I interferon signaling. Cell Metab 20, 813–826 (2014).

47. Garcia-Bermudez, J., et al. Adaptive Stimulation of Macropinocytosis Overcomes Aspartate Limitation in Cancer Cells under Hypoxia. http://biorxiv.org/lookup/doi/10.1101/2021.02.02.429407 (2021) doi:10.1101/2021.02.02.429407.

48. De Leenheer, A. P., De Bevere, V. O., Cruyl, A. A. & Claeys, A. E. Determination of serum alpha-tocopherol (vitamin E) by high-performance liquid chromatography. Clin Chem 24, 585–590 (1978).

49. Mathieu, R. E. & Riley, C. P. Quantitation of Ubiquinone (Coenzyme Q₁₀) in Serum/Plasma Using Liquid Chromatography Electrospray Tandem Mass Spectrometry (ESI-LC-MS/MS). Methods Mol Biol 1378, 61–69 (2016).

50. Lawrence, R. et al. Evolutionary differences in glycosaminoglycan fine structure detected by quantitative glycan reductive isotope labeling. J Biol Chem 283, 33674–33684 (2008).

51. Basu, A. & Weiss, R. J. Glycosaminoglycan Analysis: Purification, Structural Profiling, and GAG-Protein Interactions. Methods Mol Biol 2597, 159–176 (2023).

52. Bezwada, D. et al. Mitochondrial metabolism in primary and metastatic human kidney cancers. bioRxiv 2023.02.06.527285 (2023) doi:10.1101/2023.02.06.527285.

53. Elias, R. et al. A renal cell carcinoma tumorgraft platform to advance precision medicine. Cell Rep 37, 110055 (2021).

54. Pavía-Jiménez, A., Tcheuyap, V. T. & Brugarolas, J. Establishing a human renal cell carcinoma tumorgraft platform for preclinical drug testing. Nat Protoc 9, 1848–1859 (2014).

55. Quintana, E. et al. Efficient tumour formation by single human melanoma cells. Nature 456, 593–598 (2008).

